# Constricted migration is associated with stable 3D genome structure differences in cancer cell

**DOI:** 10.1101/856583

**Authors:** Rosela Golloshi, Christopher Playter, Trevor F. Freeman, Priyojit Das, Thomas Isaac Raines, Joshua H. Garretson, Delaney Thurston, Rachel Patton McCord

**Affiliations:** Biochemistry & Cellular and Molecular Biology Department, University of Tennessee, Knoxville, TN; UT-ORNL Graduate School of Genome Science and Technology, University of Tennessee, Knoxville, TN

**Keywords:** Constricted migration, Hi-C, 3D genome, melanoma, genome compartments, metastasis, heterochromatin, lamin, nucleus structure

## Abstract

To spread from a localized tumor, metastatic cancer cells must squeeze through constrictions that cause major nuclear deformations. Since chromosome structure affects nucleus stiffness, gene regulation and DNA repair, here we investigate the relationship between 3D genome structure and constricted migration in cancer cells. Using melanoma (A375) cells, we identify phenotypic differences in cells that have undergone multiple rounds of constricted migration. These cells display a stably higher migration efficiency, elongated morphology, and differences in the distribution of Lamin A/C and heterochromatin. Hi-C experiments reveal differences in chromosome spatial compartmentalization specific to cells that have passed through constrictions and related alterations in expression of genes associated with migration and metastasis. Certain features of the 3D genome structure changes, such as a loss of B compartment interaction strength, are consistently observed after constricted migration in clonal populations of A375 cells and in MDA-MB-231 breast cancer cells. Our observations suggest that consistent types of chromosome structure changes are induced or selected by passage through constrictions and that these may epigenetically encode stable differences in gene expression and cellular migration phenotype.

## Introduction

Despite significant improvements in the diagnosis and treatment of cancer, most patients with advanced metastatic disease face a terminal illness incurable by current therapeutic methods. The dissemination of cancer cells from the primary tumor to colonize to distant sites involves a complex multi-step invasion-metastasis cascade (vChambers *et al*, 2002; Fidler, 2003; Gupta & Massague, 2006; Lambert *et al*, 2017). To metastasize, cancer cells must squeeze through constrictions in the extracellular matrix or endothelial lining that are much smaller than their nucleus, causing major nuclear deformations. While the cell membrane and cytoplasm are quite elastic, the ability of the cell’s nucleus to withstand large deformations and squeeze through these small spaces is limited by its size and stiffness, posing challenges to constricted migration (Davidson *et al*, 2014; Denais *et al*, 2016; Friedl *et al*, 2011; Fu *et al*, 2012; Irianto *et al*, 2017c; Raab *et al*, 2016; Wolf *et al*, 2013)

Nuclear stiffness and deformability depend on two major components: lamin proteins (Lamin A/C and Lamin B) and chromatin state (Davidson *et al*., 2014; Harada *et al*, 2014; Stephens *et al*, 2017). Lamin A/C expression and its stoichiometric relation to Lamin B contributes to nuclear mechanical properties (Harada *et al*., 2014; Lammerding *et al*, 2006; Swift *et al*, 2013). Decreased expression of Lamin A/C facilitates constricted migration of mouse embryonic fibroblasts (MEFs) while increased Lamin A/C expression inhibits constricted migration of neutrophils (Davidson *et al*., 2014; Rowat *et al*, 2013). Similarly, low levels of Lamin A/C are correlated with increased aggressiveness in some cancers (Broers *et al*, 1993; Wazir *et al*, 2013), though this is not always the case (Venables *et al*, 2001; Willis *et al*, 2008). Chromatin state has also been shown to influence nuclear physical properties: increasing heterochromatin increases nuclear stiffness while increasing euchromatin decreases stiffness and increases nuclear deformability (Stephens *et al*., 2017; Stephens *et al*, 2018). Such global changes in chromatin state can also influence constricted migration. For example, increasing euchromatin or decreasing heterochromatin inhibits melanoma and fibrosarcoma cell migration (Gerlitz & Bustin, 2010; Krause *et al*, 2019; Maizels *et al*, 2017; Panagiotakopoulou *et al*, 2016; Segal *et al*, 2018). Chromatin decondensation was found to increase breast cancer cell invasion into dense extracellular matrices but decrease invasion into loose matrices (Fischer *et al*, 2020). The loss of SETDB1 methyltransferase in lung cancer leads to reorganization of heterochromatin and associates with changes in migration (Zakharova *et al*, 2021). Extracellular physical forces can also affect nuclear mechanics and chromosome structure. Mechanical cues in a cell’s environment can lead to a spatial redistribution of chromatin (Heo *et al*, 2021). Irreversible stretching of chromatin domains has been observed in some nuclei aspirated into narrow micropipettes (Irianto *et al*, 2017b), and external cellular forces can even affect gene expression through direct chromatin deformations (Tajik *et al*, 2016). Despite such connections between chromatin structure, nucleus deformation, and nucleus mechanics, it is not fully understood whether the 3D organization of the genome affects or is affected by constricted migration.

Ever-improving microscopic techniques and the development of chromosome conformation capture approaches have revolutionized our understanding of 3D genome architecture in the past decade (Abbas *et al*, 2019; Bickmore & van Steensel, 2013; Dekker *et al*, 2002; Pombo & Dillon, 2015; Rowley & Corces, 2018). Chromosomes are folded at different length scales into loops, topologically associating domains (TADs), active and inactive (A/B) compartments, and chromosomal territories. How these layers of structure are affected by physical nucleus shape changes such as the kind experienced during constricted migration remains largely unknown. Previous work has shown that 3D chromosome organization changes correlate with the progression of cancer (Barutcu *et al*, 2015; Taberlay *et al*, 2016; Zhou *et al*, 2019) and that both the formation and downstream effect of chromosomal translocations in cancer progression is influenced by 3D genome structure (Hnisz *et al*, 2016; Zhang *et al*, 2012). In addition, some changes have been noted in the 3D genome structures of neutrophils that have undergone constricted migration (Jacobson *et al*, 2018) and there is evidence that the process of constricted migration itself could alter chromatin modifications (Hsia *et al*, 2021). Taken together, previous evidence suggests that chromosome structure may influence a cancer cell’s ability to undergo constricted migration and may be influenced by the constriction itself. In this study, we seek to unite these previous ideas and investigate what 3D genome structures of cancer cells are associated with constricted migration and how these structures relate to gene regulation and cancer cell phenotype.

Here, we use invasive human melanoma cells (A375) to investigate the properties of the nucleus and 3D chromosome structure that accompany constricted migration. We find that A375 cells that have migrated numerous times through tight constrictions show increased migration efficiency and are phenotypically distinct from cells that do not pass through constrictions. These repeatedly constricted cells exhibit specific and stable differences in 3D genome organization, particularly at the level of A/B compartmentalization, Lamin A/C localization, and gene expression patterns. We observe similar trends of 3D genome structure changes across multiple different clonal populations of A375 cells. Some features of these structural changes are conserved in MDA-MB-231 breast cancer cells that have undergone constricted migration. Our results suggest that 3D genome architecture correlates with the ability of cancer cells to undergo constricted migration. Our experimental data and modeling suggest that 3D genome and nucleus differences could arise from a combination of selection from an initially heterogeneous cell population and changes induced by constricted migration itself.

## Results

### Melanoma cells exhibit an increased migration efficiency and differences in cell and nuclear morphology after multiple rounds of constricted migration

Previous work has shown that passage through sequential rounds of constricted migration can lead to cellular and genomic alterations (Irianto *et al*, 2017a). To investigate potential nucleus and 3D genome alterations associated with constricted migration, we allowed A375 cells to undergo 10 rounds of migration through 5μm Transwell filters (Figure 1A, Methods). The minimum diameter of A375 nuclei ranges from 8-16μm (Figure S1A), so passing through these 5μm pores requires nuclear constriction. Cells that successfully migrated through 5 µm constrictions each round were more efficient at migrating in subsequent rounds of the same Transwell assay, reaching 70% migration after 10 consecutive rounds (Bottom-5 cells) (Figure 1B and Figure S1B). In contrast, A375 cells that failed to undergo migration through 5 μm pores showed a progressive decrease in migratory efficiency with each round, approaching 0% migration after 10 consecutive rounds (Top-5) (Figures 1B and S1B). These differences in migration efficiency correspond to other notable differences in cell phenotype. Phase contrast imaging showed that Bottom-5 cells exhibit fewer cell-cell adhesions and an elongated cell body when compared to Top-5 cells (Figures 1B, S1E). Live cell imaging of Top-5 and Bottom-5 cells migrating on a 2D surface showed that Bottom-5 cells display elongated shapes and frequent extension of protrusions while Top-5 cells maintained more rounded shapes with ruffled leading edges during a 13-hr migration period (Figure S1G and Movies S1 and S2). The migration efficiency differences of Bottom-5 cells were maintained even after continuous culture (5 passages) or freezing and thawing, indicating that this highly migratory phenotype is stable (Figure S1H). Similar cellular phenotypes were observed in a sequential constricted migration replicate extended to 20 rounds of migration (Bottom20-5, Figure S1C-E).

**Figure 1.**
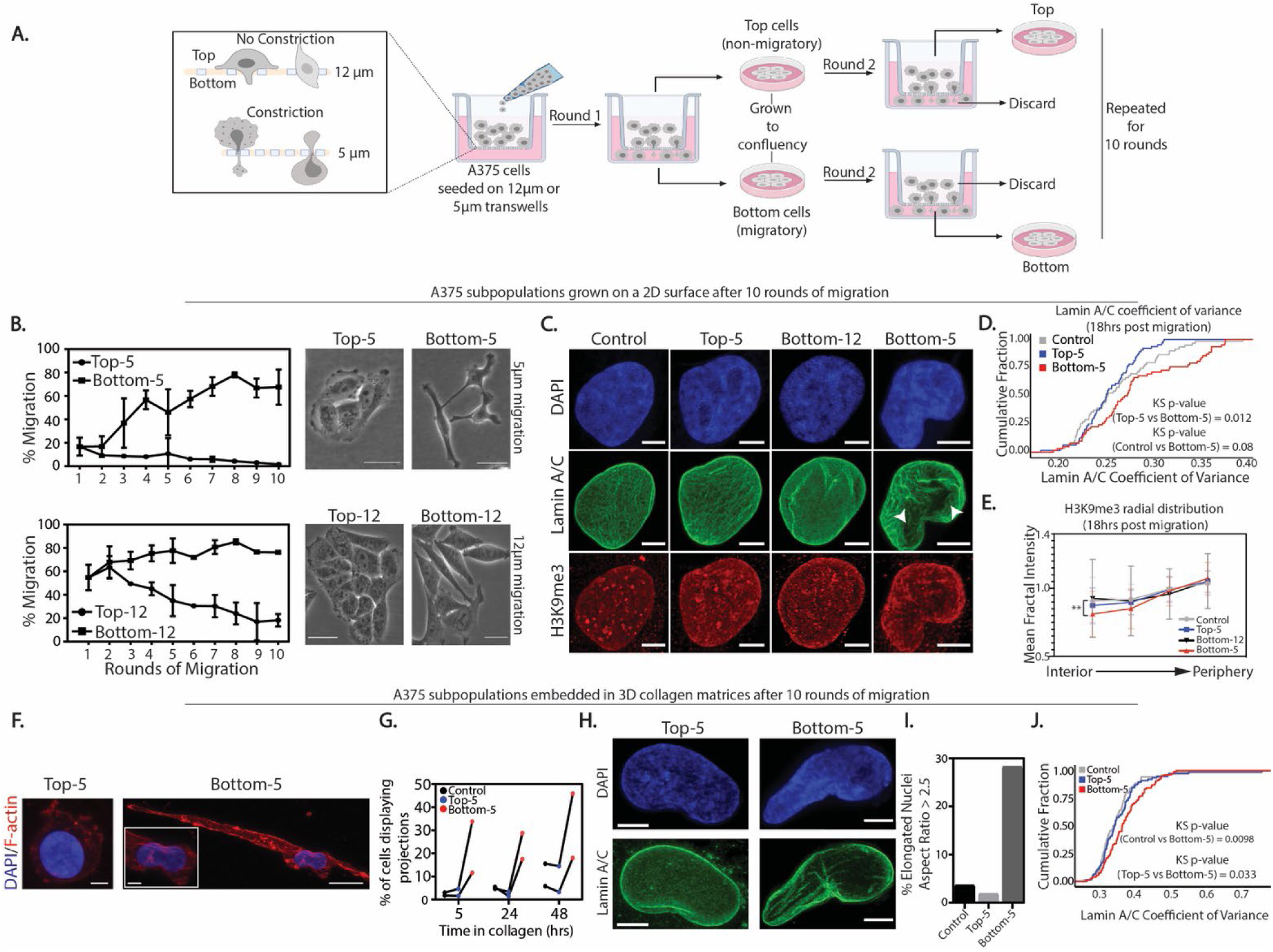
A375 cells exhibit morphological differences and increased migration efficiency after sequential rounds of constricted migration. (A) Layout of sequential Transwell migration experiment with A375 cells. (B) Migration efficiency (% of cells that migrated through filter pores in each round) of A375 cells during 5 μm constricted migration and 12 μm migration (left panels). Error bars = mean ± SD for 3 biological replicates. Phase contrast imaging of the final cell populations after the 10 rounds of migration, cells cultured on 2D plastic (right panels, Scale bar = 50 µm). (C) A375 nuclei stained with DAPI (blue), Lamin A/C (green) and H3K9me3 (red) in the indicated migration conditions. Images shown represent a maximum projection of z-stacks (Scale bar= 5 µm). (D) Cumulative distribution plot of Lamin A/C coefficient of variance in all indicated A375 subpopulations (Control (n = 59), Top-5 (n = 64), Bottom-5 (n = 62) nuclei). Kolmogorov-Smirnov test p-values indicated in plot (E) Radial intensity distribution of H3K9me3 signal in the maximum projected nuclei of Control (n = 59), Top-5 (n = 63), Bottom-5 (n = 64) and Bottom-12 (n = 80) nuclei (** p = 0.0069, two-tailed t-test). Error bars = mean ± SD. (F) Maximum projection confocal images of Top-5 and Bottom-5 cells embedded in collagen and stained with DAPI (blue) and Phalloidin (red). (Inset) Zoom in of nucleus of Bottom-5 cell. Top-5 and Bottom-5 zoomed inset scale = 5 μm; Bottom-5 zoomed out image scale = 20 μm. (G) Fraction of Control, Top-5 and Bottom-5 cells displaying projections at indicated timepoints after cells were embedded in 3D collagen matrices. Each biological replicate is connected by a line. (H) Nuclei of Top-5 (left) and Bottom-5 cells (right) embedded in 3D collagen matrix and stained with DAPI (blue) and Lamin A/C (green). Scale = 5μm. For 3D reconstruction visualization, see Movies S5 and S6. (I) Percent of cells embedded in collagen that had elongated nuclei (Aspect Ratio > 2.5) for A375 Control (n = 54), Top-5 (n = 52) and Bottom-5 (n = 53). (J) Cumulative distribution plot of Lamin A/C coefficient of variance for Control (n = 70), Top-5 (n = 78) and Bottom-5 (n = 109) cells embedded in 3D collagen matrices. Kolmogorov-Smirnov test p-values indicated in plot.

To investigate whether the differences we observed were specific to passing through a constriction, we repeated the sequential migration experiment using Transwell filters with pores larger than the majority of A375 nuclei (12 µm). As expected, A375 cells migrated at a much higher rate (∼ 55-70%) in the first round of this large pore Transwell assay (Figures 1B and S1I). Cells that were successful at migrating through 12 µm pores for 10 rounds (Bottom-12) displayed only a minimal increase in migratory efficiency. In contrast, the progressive decrease in migration efficiency of cells that did not pass through the 12 µm pores in each round (Top-12) suggests that there exists a sub-population of cells that are generally poor at migration (Figures 1B and S1I). While Bottom-12 cells displayed some loss of cell-cell adhesion, quantification of phase contrast images revealed no statistically significant cell morphology differences between Top-12 and Bottom-12 cells (Figure 1B and S1J-K). When challenged to migrate through 5 μm pores, Bottom-12 cells showed a migration efficiency similar to the initial control population (∼ 3-20 %). These results indicate that simply being able to migrate is not enough to guarantee efficient migration through constricted spaces, indicating that additional features are associated with constricted migration.

Given the large nucleus deformations necessary for constricted migration, we next turned to investigating the nuclear architecture of sequentially migrated A375 cells. We evaluated nucleus morphology and the distributions of Lamin A/C and heterochromatin (H3K9me3) 18 hours after sequential constricted migration using immunofluorescence imaging (Figure 1C). We found that sequentially constricted Bottom-5 cells showed a tendency to have more elongated nuclei compared with control cells or unconstricted Bottom-12 cells (Figure S2A,B). No significant differences in the overall intensity of Lamin A/C were observed in Bottom-5 cells 18 hours after constricted migration (Figure S2C). Earlier timepoints after constricted migration suggested a transient decrease in Lamin A/C intensity (Figure S2E), but this difference is not stable. Western blots confirmed the lack of change in Lamin A/C protein levels and detected no change in Lamin A/C phosphorylation at Ser22, a modification that can influence the lamina vs. nucleoplasmic localization of Lamin A/C (Torvaldson *et al*, 2015) (Figure S2I,K). Similarly, we found no global change in levels of Lamin B1 or the H3K9me3 heterochromatin mark (Figure S2J,L). These unchanged Lamin A/C levels echo recent observations reported in mechanically stretched cells (Nava *et al*, 2020) but are in contrast to other reports that show an association between lower levels of Lamin A/C and increased constricted migration ability (Harada *et al*., 2014).

Despite the lack of significant changes in overall Lamin A/C levels, we did observe a stable difference in the distribution of Lamin A/C within the nuclear lamina after constricted migration. A375 Control, Top-5 and Bottom-12 nuclei showed a fairly uniform distribution of Lamin A/C throughout the nuclear envelope, while Bottom-5 nuclei displayed areas of stronger Lamin A/C enrichment or depletion (Figure 1C, Movies S3 and S4). By visual inspection, we found that ∼50% of Bottom-5 nuclei show an abnormal distribution of Lamin A/C compared to only ∼25 % of Bottom-12 nuclei (Figure S2E) suggesting that differences in Lamin A/C distribution are specific to cells that have undergone constricted migration. Quantitatively, we found that Bottom-5 nuclei exhibit a higher Lamin A/C coefficient of variance when compared to Top-5 (Figure 1D). We also observed an increase in nuclear envelope invaginations and wrinkling in Bottom-5 nuclei (Figure S2F). Line scans across central slices of nuclei also suggested that Bottom-5 cells have a lower amount of Lamin A/C along their minor axis (narrower dimension) than Top-5 cells (Figure S2G). These alterations in lamin distribution are reminiscent of previously described redistribution of lamins as a consequence of nuclear deformation (Pfeifer *et al*, 2018).

Like Lamin A/C, we found that the distribution of heterochromatin is altered in A375 cells that have undergone constricted migration. Heterochromatic foci, as measured by H3K9me3 immunofluorescence, were distributed throughout the nucleus in Control, Top-5, and Bottom-12 cells while a larger proportion of H3K9me3 signal was localized to the nuclear periphery in Bottom-5 cells (Figure 1C, 1E, and S2G). Since heterochromatin can be tethered to the nuclear lamina through Lamin A/C (Manzo *et al*, 2022; Solovei *et al*, 2013), we evaluated whether regions of altered Lamin A/C distribution related to heterochromatin distribution. Indeed, the H3K9me3 signal was correlated with stripes of increased Lamin A/C intensity in Bottom-5 cells (Figure S2H).

When embedded in 3D collagen matrices, Bottom-5 cells displayed a higher number of projections (Figure 1F and G) accompanied by more nuclear deformations than Control or Top-5 cells (Figure 1H and I, Movies S7 and S8). Live cell imaging in collagen showed that Bottom-5 cells were more likely to invade through the collagen while Top-5 cells remained more stationary (Movies S5 and S6). In this 3D collagen context, Bottom-5 cells also showed a more variable Lamin A/C distribution (Figure 1J).

Overall, our image analyses imply that, in A375 cells, constricted migration is associated with alterations in the distribution of Lamin A/C and heterochromatin that are visible at even 18 hours after migration. Additionally, quantification of nuclear shape differences in collagen distinguished cells that had undergone constricted migration in Transwell chambers from cells that did not, indicating that the ability to deform their nucleus and migrate in a 3D environment is a key distinguishing factor among the A375 cell subpopulations.

To evaluate the impact of cancer cell type on our constricted migration observations, we repeated the sequential migration and imaging experiments in MDA-MB-231 breast cancer cells (Figure S3). These cells exhibited a higher initial ability to migrate through 5μm constrictions (∼35%) when compared to A375 cells. MDA-MB-231 cells that migrated through 5μm pores for 10 rounds (Bottom-5) showed an increase in migration efficiency, like A375 cells, though the change was not as dramatic (Figure S3A, left panel).

When challenged to migrate through large 12 μm pores, MDA-MB-231 Top-12 and Bottom-12 cells showed no significant separation in their migratory efficiency (Figure S3A, right panel), in contrast to A375 cells, which had a subpopulation that could not migrate through even large pores. This indicates that there is less initial heterogeneity among the MDA-MB-231 cells in their ability to migrate.

Indeed, whether or not they have undergone constricted migration, MDA-MB-231 cells exhibited elongated cell bodies and low cell-cell adhesions (Figure S3B). However, like A375 cells, a higher fraction of MDA-MB-231 cells that underwent constricted migration for 10 rounds (Bottom-5) showed elongated cell bodies as measured by aspect ratio (Figure S3C). MDA-MB-231 cells tend to have more elongated nuclei than A375 cells overall, but we still observed a further increase in nucleus elongation among Bottom-5 MDA-MB-231 cells (Figure S3D-E). These observations coincide with previous reports of elongated cell and nuclear shape correlated with higher migration propensity in MDA-MB-231 cells (Baskaran *et al*, 2020). As in A375 cells, the increased migratory efficiency of MDA-MB-231 Bottom-5 cells was not related to a change in overall Lamin A/C or H3K9me3 protein levels (Figure S3H-I). Differences in Lamin A/C distribution in MDA-MB-231 cells were more subtle than in A375 cells. No significant differences were observed among Control, Top-5 and Bottom-5 cells in Lamin A/C intensity variance (Figure S3F), but a higher fraction (90%) of Bottom-5 cells displayed visibly wrinkled nuclei compared to Control and Top-5 (75-80%, Figure S3G). Like A375 cells, MDA-MB-231 Bottom-5 cells have a higher H3K9me3 intensity at the nuclear periphery across the major and minor axis when compared to Control and Top-5 cells (Figure S3J).

In both these cancer cell types, constricted sequential migration can result in a more highly migratory subpopulation that exhibits some nucleus morphology changes and differs in the distribution, but not the levels of Lamin A/C and H3K9me3. Given the role of lamins and heterochromatin in organizing the 3D genome structure, we next asked whether differences are found in the 3D organization of chromosomes in sequentially constricted cells.

### Chromosome compartment changes observed after rounds of constricted migration

To measure changes in chromosomal contacts in A375 cells, we performed genome-wide chromosome conformation capture (Hi-C) on all subpopulations of sequential constricted and unconstricted migration conditions (Methods, Table S1). Strikingly, using 250 kb binned matrices, we observed regions of the genome that had a different pattern of interactions in Bottom-5 cells compared to all other conditions (Figure 2A, example indicated by black arrows). Since these changes primarily affected the plaid pattern that indicates spatial compartmentalization, we performed Principal Component Analysis (PCA) to classify regions according to their A/B compartment identity. We identified regions of the genome that have a consistent compartment identity among all unconstricted conditions (Control, Top-12, Top-5, Bottom-12) but have a different compartment identity in Bottom-5 cells (Figure 2B, boxed region, for example). At a stringent threshold (see Methods) we observed that 2.89% of 250 kb regions across the genome switched from the B (typically more heterochromatic) to A (typically more euchromatic) compartment and 1.63% of the bins switched from the A to B compartment in sequentially constricted Bottom-5 cells compared to Control (Figure 2C, middle panel). Cells that migrated through large pores (Bottom-12) show a minor shift in the compartment eigenvector in some of these regions (Figure 2B and S4D), but overall have 10-fold fewer compartment switches than constricted cells (Figure 2C, left panel). These compartment switches are reproduced in replicates, including an independent experiment in which we extended migration through 5 µm pores to 20 rounds (Bottom20-5) (Figure S4B-C). Some regions (0.5% of bins) had a different compartment identity in cells that were unable to migrate even through large pores (Top-12) indicating that there may be a distinct epigenetic structure of non-migratory cells as well (Figure S4A). Overall, genome-wide hierarchical clustering analysis of the compartment eigenvectors segregates conditions according to whether they have undergone constricted migration (Figure 2D).

**Figure 2.**
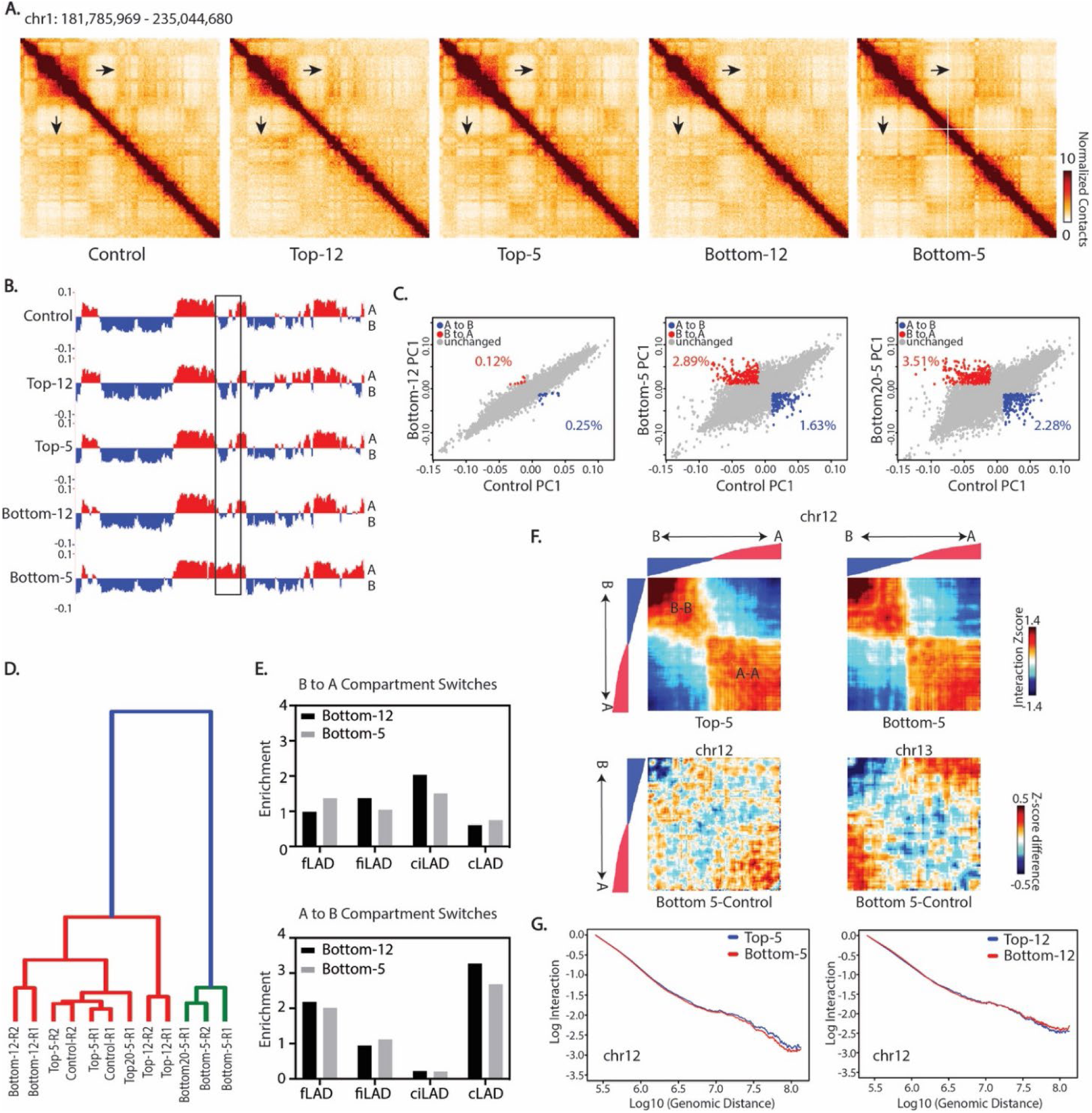
Compartment identity switches after constricted migration. (A) 250 kb binned Hi-C interaction heatmaps of chr1 (chr1:181,785,969 - 235,044,680) across A375 conditions. Arrows: locations of visible difference in Bottom-5 contact maps. (B) PC1 track of compartment identity (250 kb bins) in the same region of chr1. Boxed region highlights the same compartment switch highlighted by arrows in A. (C) Compartment PC1 value comparisons for each bin genome wide in Bottom-12 (left), Bottom-5 (middle) and Bottom20-5 (right) vs. Control. Percentage of bins that met criteria for “compartment switch” from B to A (red; PC1 < -0.01 to > 0.01) or A to B (blue; PC1 > 0.01 to < -0.01) indicated on each panel. (D) Clustering of all A375 conditions by genome-wide compartment PC1. (E) Enrichment of different LAD types in regions that switch their compartments from B to A and A to B in Bottom-5 and Bottom-12 cells. (F) Above: chr12 saddle plots for Top-5 and Bottom-5 conditions show interaction Z-score (normalized by generic distance decay) between all bin pairs that do not switch compartments, sorted by Control compartment PC1 value from strongest B-B to strongest A-A. Below: change in interaction Z-score from Control to Bottom-5 conditions for chr12 (left) and chr13 (right). Track above heatmaps = ordered PC1 values of A375 Control cells. (G) Whole genome contact scaling plots over distance for chr12 at 250 kb bin size for both 5 µm and 12 µm Transwell migration.

Since compartmentalization involves the spatial segregation of heterochromatin and euchromatin, these changes may relate to our microscopic observations of increased H3K9me3 peripheral localization after constricted migration (Figure 1). To investigate this link further, we quantified whether genomic regions that change compartment identity in Bottom-5 cells correspond to certain lamin associated domain (LAD) types, as described in a previously established LAD atlas across a cohort of nine different cell types (Carolyn de Graaf, 2019; Kind *et al*, 2015). We observed that while B to A switches were fairly evenly distributed across LAD types, A to B switches tended to occur in facultative LADs (fLADs and fiLADs; regions that vary in their lamin association across cell types) and constitutive LADs (cLADs; regions found at the lamina in most cell types) (Figure 2E). This suggests a possible further spatial consolidation of heterochromatin regions corresponding to the observed coalescence of H3K9me3 at the nuclear periphery.

Just as the cellular morphology changes in MDA-MB-231 cells after constricted migration are less dramatic than in A375 cells, this cell type also shows fewer compartment switches with sequentially constriction than the A375 cells did (Figure S5A). The initial genome structures of these different cancer cell lines are quite different, so we would not expect compartment changes to occur in the exact same locations. However, we do observe 79 bins that switch from A to B and 36 bins that switch from B to A in both A375 and MDA-MB-231 constricted migration (Figure S5B). Interestingly, some of these commonly switched regions contain genes relevant to migration related phenomena, including extracellular matrix organization, actin filament polymerization, and chemotaxis (Figure S5C).

### Weaker B compartment interactions and increased intercompartmental interactions are a conserved feature of constricted cells

While the switching of compartment identity after constricted migration is a striking phenomenon, and clearly apparent by visual inspection of the contact maps, only ∼ 4.5% of the genomic bins were affected by a complete compartment switch in A375 cells according to our thresholds. To evaluate how constricted migration may impact the rest of 3D genome, we evaluated contacts between regions not involved in compartment identity switches using “saddle plots”, ordering regions from strongest B to strongest A bins based on PC1 values (Imakaev *et al*, 2012). We observed an evident contact separation between the B and A compartment both with and without constricted migration (Figure 2F). However, the relative strength of inter and intra-compartment contacts changed in migrated cells. Bottom-5 cells exhibited a loss in the strongest B-B interactions and an overall increased mixing of the strongest B and strongest A compartments (Figure 2F, bottom panels). In unconstricted migration, we see apparent overall loss of compartment strength (Figure S4D), though this effect on both compartments could also be explained by the overall higher background noise in this condition.

In contrast to these changes in interaction strength within and between compartments, we overall observed negligible changes in the chromosome contact scaling with distance after migration (Figure 2G). This interaction scaling reflects local chromatin compaction and average loop size and density. Further, we observed no clear differences between TAD boundary strength in any of the conditions (Figure S4E). These observations suggest that the major differences with constricted migration occur in larger scale rather than smaller scale chromosome structures and are in line with previous reports that compartment-scale interactions can change independently from TAD-scale contacts (Nuebler *et al*, 2018).

Interestingly, we saw a similar compartment strength effect in MDA-MB-231 cells that have undergone sequential migration through constriction. MDA-MB-231 Bottom-5 cells lose strong B compartment interactions while gaining interactions between the A and B compartment (Figure S5D). This loss of interactions in the B compartment was also previously reported in neutrophils after passage through small pores, suggesting it may be a general feature of constricted migration chromosome structural changes (Jacobson *et al*., 2018).

### Gene expression changes after constricted migration reflect migratory potential

Since spatial compartmentalization generally correlates with gene expression, we next investigated potential differences in gene expression in cells that have undergone constricted migration. We performed RNA-Seq on all A375 subpopulations. We identified 977 and 1473 genes that were differentially expressed in Bottom-5 and Bottom20-5 relative to Control, respectively (Table S2). Hierarchical clustering of all conditions, starting with the subset of genes differentially expressed in Bottom-5 or Bottom20-5, segregated cells that undergo constricted migration from all other conditions (Figure 3A and Figure S6A). The 482 genes that were upregulated in Bottom-5 and Bottom20-5 and downregulated in other conditions are enriched in pathways related to metastasis, such as TGF-beta, TNF-alpha, EGFR and Integrin β-4 signaling pathways (Figure 3B). On the other hand, the 368 genes downregulated in Bottom-5 and Bottom20-5 but upregulated in other conditions are enriched in developmental pathways. Since melanoma cells originate from the neural crest, this cluster may relate to downregulation of melanocyte-specific gene expression in constricted migrating cells (Figure 3B). We also observe downregulation of a group of genes important for cell-cell/matrix adhesion (CADM3, CADM1, ITGA9). A third cluster of genes, downregulated in Bottom20-5 and upregulated in Top-5 are involved in cell adhesion, which correlates with the differences we observed in cell-cell contacts between these subsets (Figure 1C).

**Figure 3.**
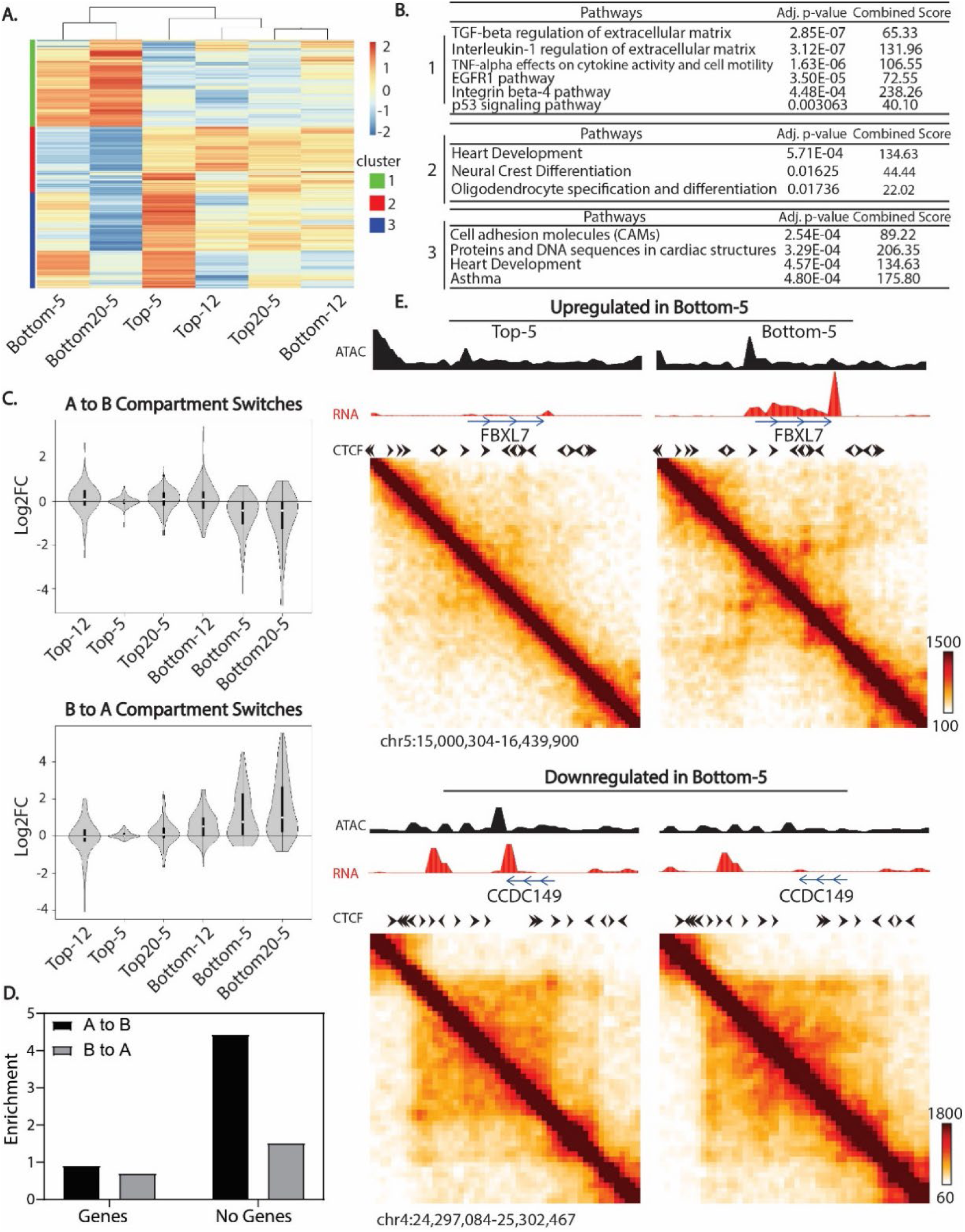
Changes in gene expression correlate with metastatic ability and chromosome structure changes. (A) Hierarchical clustering of RNA-Seq datasets based on differentially expressed genes of Bottom-20 cells relative to Control (Log2 fold change >1). Color scale indicates log2(Condition/Control) expression for each gene. (B) Pathway enrichment analysis of identified gene clusters from (A) using BioPlanet 2019 package from Enrichr. (C) Log2FC of expression in the indicated condition vs. Control for genes located in regions that switch compartments from A to B (top panel) and B to A (bottom panel) in Bottom-5 cells. (D) Bins involved in compartment switches were classified as either containing or not containing annotated transcripts and then the Observed fraction in each category was compared to the Expected representation in a randomly selected set. (E) Contact maps of regions around genes upregulated (FBXL7) and downregulated (CCDC149) in Bottom-5 cells. Tracks over the matrices represent ATAC-Seq (black) and RNA-Seq (red). The tracks have been smoothed with a 40 kb sliding window, 20 kb step. Black arrows indicate known CTCF motif directionality.

When we specifically compared constricted vs. unconstricted migration (Bottom-5 vs Bottom-12), we found genes more highly expressed in constricted cells are related to focal adhesion pathways and regulation of the actin cytoskeleton (Figure S6B). These changes correlated with the increase in protrusions and changes in cell shape observed for Bottom-5 cells in Figure 1 and S1. We also observed a differential gene expression signature for Top12 cells, which did not migrate even through large pores, relative to the rest of the A375 population. The majority of genes downregulated in Top-12 cells were related to TGF-beta, EGFR, integrin and p-53 signaling, all pathways related to cancer cell migration and metastasis (Figure S6B).

These gene expression changes correlate with compartment changes observed upon constricted migration. Genes found in regions that switched their compartment identity from A to B in Bottom-5 cells displayed an overall decrease in expression while genes in regions that switched from B to A tended to show increased expression when compared to other conditions (Figure 3C). Even in regions that did not switch compartments, we found that changes in interaction strength within compartments also correlated with changes in gene expression in Bottom-5 cells (Figure S6D). Genes in regions that exhibited B-B interaction loss in Bottom-5 cells showed an overall increase in expression in these cells. These observations suggest that some of the compartment changes associated with constricted migration directly relate to changes in gene regulation.

Gene expression changes do not account for all compartment changes, however. Notably, we found that among the genomic bins that switched from A to B and B to A compartment, a larger than expected fraction contain no genes (4-fold and 2-fold enrichment over random expectation, respectively) (Figure 3D). This observation suggests that some structural changes after migration are not directly related to gene regulation changes and might relate to mechanical properties of the nucleus.

We next examined whether observed gene expression changes relate to local genome structure changes. At highly upregulated genes such as the FBXL7 gene (∼ 8-fold upregulated in Bottom-5 and Bottom20-5), we observed the emergence of a domain-like structure in Bottom-5 cells and an increase in chromatin accessibility (Figure 3E, top panel). Conversely, genes that were downregulated after constricted migration often showed a loss of loop structures and chromatin accessibility (SOD3 and CCDC149 3-15 fold downregulated; Figure 3E, bottom panel). However, we also encountered cases in which the genes were differentially expressed such as PLEKHA6 (log2FC > 2 in Bottom-5 and Bottom20-5) and PKDCC (log2FC = -2.5 in Bottom-5 and Bottom-20) but their local genomic structures remained similar with modest changes in chromatin accessibility (Figure S6C). The loss or formation of loops or domains associated with changes in transcription occurred in regions that had few or no other strong architectural loops, such as the kind mediated by CTCF. Genes that changed expression without notable 3D looping changes, in contrast, were located in regions with many other loops and TAD boundaries. These observations support the idea that transcription driven domains can arise separately from CTCF-based domains (Rowley *et al*, 2017; Zhang *et al*, 2020). Further, the transcription-associated CCDC149 loop occurs between an existing TAD boundary and a promoter, and could be explained by a collision between opposing direction loop extrusion and RNA polymerase activity, as suggested in other systems (Brandao *et al*, 2019).

Previous work has observed gene expression changes induced by mechanical stress, such as stretching (Le *et al*, 2016; Nava *et al*., 2020). We observed little correlation between differentially expressed genes in cells that have undergone constricted migration and cells that have been mechanically stretched (Figure S6E). This points to differences between a cell being passively deformed and one that is actively coordinating its own migration through a tight constriction.

### Altered translocations and copy number variation after constricted migration

Constricted migration has previously been shown to result in DNA damage and genome instability (Irianto *et al*., 2017a). Since Hi-C can detect structural aberrations such as translocations, we examined interchromosomal interactions in 2.5 Mb binned matrices to search for potential genomic aberrations after constricted migration (Figure 4).

**Figure 4.**
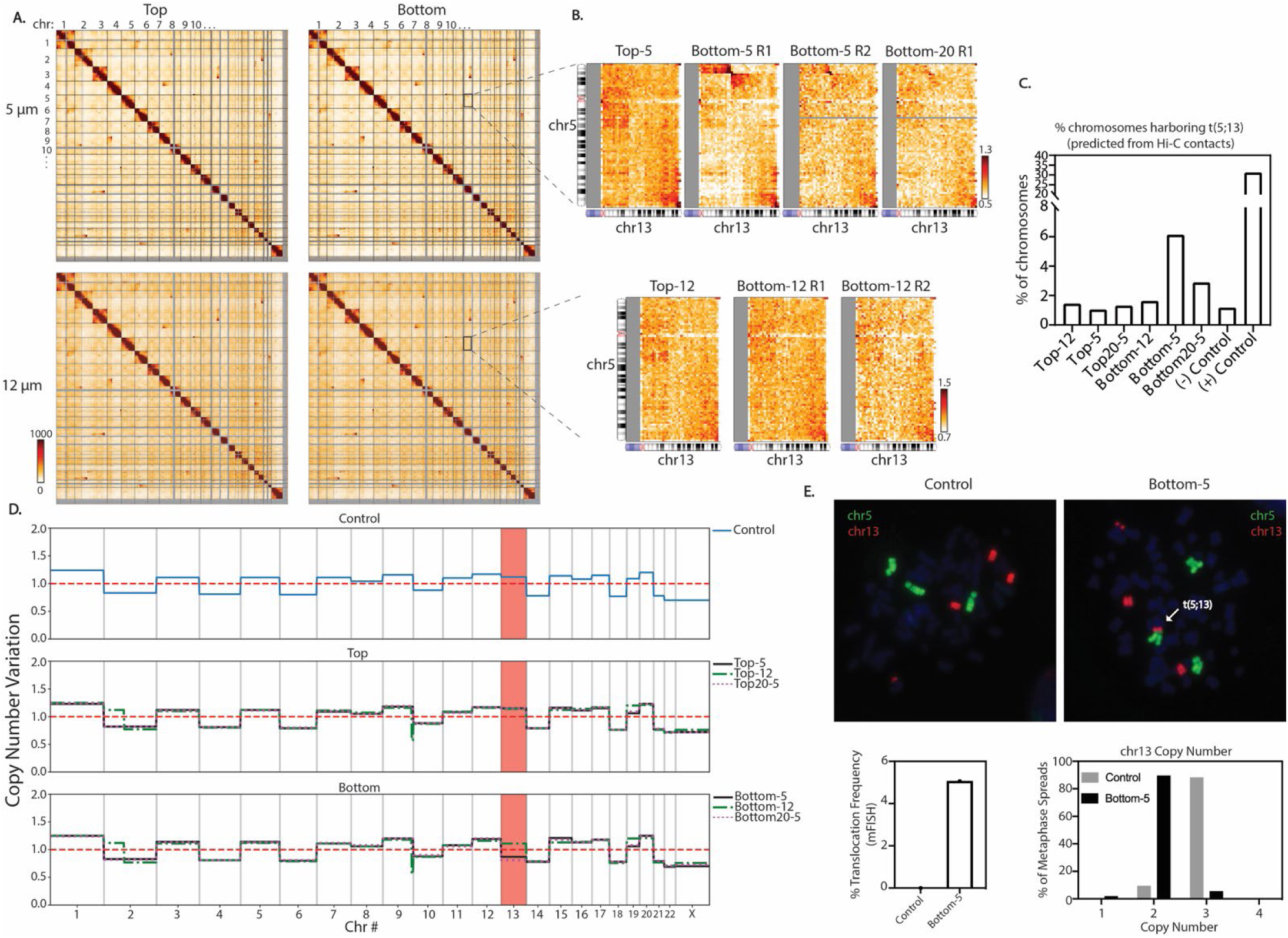
Global genomic rearrangements after constricted migration. (A) Whole genome 2.5 Mb Hi-C interaction frequency maps for cells that have undergone 5 and 12 μm Transwell migration. (B) Zoom-in of inter-chromosomal interactions between chr5 and chr13 for cells that have undergone 5 and 12μm Transwell migration. All biological replicates performed are shown for Bottom-5 and Bottom-12. (C). Estimation of the % of chromosomes bearing the t(5;13) based on the frequency of the proposed translocation interaction divided by the average neighboring bin cis interactions. (D) Copy number variation among all A375 subtypes inferred from total raw Hi-C counts. Red dashed line represents the mean copy number level while the other lines represent the copy number of A375 cell subpopulations. The region highlighted in red shows that chr13 exhibits copy number loss after 10 rounds of constricted migration. (E) Metaphase spread labeled with chromosome specific FISH probes for chr5 (green) and chr13 (red). Bottom graphs represent frequency of t(5;13) quantified by FISH (left) and copy number quantification for chr5 and chr13 (right).

Translocations inherent to A375 are present in all sub-populations of cells (Figure 4A). However, a new translocation between chr5 and chr13 (t(5;13)) was evident in each replicate of cells that had undergone constricted migration (Bottom-5 and Bottom20-5) but not unconstricted migration (Figure 4B). The specific breakpoint region of this translocation, as detected by a dramatic change in Hi-C contacts between chr5 and chr13 (Figure S7A), falls in a gene poor region of both chromosomes. We next used Hi-C contacts to estimate what fraction of the chromosomes in the population contain this translocation (see Materials and Methods for detailed analysis approach). We estimated that this translocation is present in 6.13% of chromosomes in Bottom-5 and 2.89% in Bottom-20 (Figure 4C). This translocation was undetected in cells that did not undergo constricted migration (Control, Top-12, Top-5, Top20-5 and Bottom-12): interactions at this region were indistinguishable from background interchromosomal levels in these cells (Figure 4C). Using chromosome specific FISH probes, we confirmed the presence of the t(5;13) translocation in Bottom-5 cells and found that it occurred in about 5% of cells (Figure 4E). In contrast, we never observed this translocation in the initial A375 population, even though our sample size in the FISH experiment should allow us to detect a translocation as rare as 0.25% with 95% confidence.

In addition to translocations, changes in chromosome copy number have been previously reported after constricted migration (Irianto *et al*., 2017a). We therefore followed a published approach (Servant *et al*, 2018) to use raw read counts from our Hi-C data to infer copy number variations in the constricted subpopulation of cells. The control population of A375 cells already exhibits non-uniform copy number across chromosomes and chromosome regions (Figure 4D, top panel). Most chromosomes maintained their copy number independent of constricted migration, but cells that had undergone constricted migration (Bottom-5 and Bottom20-5) showed notable copy number loss of chr13 (Figure 4D, bottom panel). This chr13 copy number loss (3 copies to 2 copies) was also confirmed by FISH (Figure 4E).

In the MDA-MB-231 breast cancer cells, we observed an even more dynamic picture of translocation changes with constricted migration (Figure S5F and G). While many of the translocations observed in the initial population are maintained, 6 translocations were lost while 6 others were gained in the cells that passed through constrictions (Figure S5G). This suggests an overall higher level of genomic instability in this cell line as compared with A375 cells. In both cell lines, these translocation changes may in part result from selection from a heterogeneous initial population or may reflect genomic instability induced by the constricted migration itself.

Beyond revealing translocations, interchromosomal contact data can reveal changes in patterns of whole chromosome territory interactions between conditions. In A375 cells, we observe a decrease in interchromosomal interactions of chr13 (Figure S7B, top panel) in cells that had passed through constrictions. This is not likely to be explained purely by copy number loss at chr13, as this would decrease both inter and intra-chromosomal contacts rather than changing the ratio between the two. Additionally, smaller chromosomes (chr14-chr22) display an increase in interactions with other chromosomes in Bottom-5 but not Bottom-12 cells (Figure S7B). This measurement could indicate a constricted-migration associated change in the relative compactness and intermixing of the smaller gene-dense chromosomes.

### Agent based modeling reveals that heterogeneity and constricted migration induced changes could both contribute to sequential migration effects

The presence of the subpopulation of A375 cells that does not migrate at all (Top-12) suggests an initial heterogeneity in migratory ability within the population, as previously reported in this cell line (Kozlowski *et al*, 1984). However, it is not clear whether the increase in migratory efficiency and associated 3D genome and nucleus morphology alterations in Bottom-5 cells are induced by repeated constricted migration or represent a pre-existing population selected by the constriction (Figure S8A). As a first step to evaluate which of these options could explain our observed migration results, we used agent-based modeling (ABM) to simulate a variety of initial population compositions and changes with migration (Figure S8B; see Materials and Methods for more detail). This model reports what fraction of cells would pass through a Transwell filter at each round of migration based on probabilistic choices of migration and division (Figure S8C). If the initial population contains a mixture of moderate and super-migrators, the migration efficiency of Top cells does not decline over the rounds of migration, which does not replicate the results of A375 cells, but does in some ways mimic the behavior of MDA-MB-231 cells (Figures S8B and S8D, first panel). The A375 experimental data can be recapitulated if the initial starting population is composed of three heterogeneous subpopulations (non-migrators, moderate migrators, and super migrators) with highly migratory cells representing 0.5% of the population (Figure S8D, second panel). If this scenario is true, then our Bottom-5 specific 3D genome and nucleus structures may be pre-existing in a small fraction of the initial population that are primed for constricted migration.

However, we are also able to recapitulate our experimental data with a scenario in which there are only non-migrators and moderate migrators in the initial population but then cells that squeeze through constrictions experience an induced change that alters their phenotype (boosting function shown in Figure S8D, third panel). In contrast, this same mixed population of non-migrators and moderate migrators without induced changes cannot recapitulate the increase in migration efficiency we observe (Figures S8D, fourth panel). Overall, our modeling results suggest that A375 cells must have an initial non-migratory subpopulation, but that the increase in migratory efficiency with constriction could either be a selection or induction process or a combination of both.

### Clonal populations of A375 cells exhibit similar trends of 3D genome structure change with constricted migration

To provide an additional experimental perspective on the relative contribution of initial population heterogeneity to the differences we observe after constricted migration, we isolated single cells from the parent A375 population (here called A375-P) and grew them into four different clonal populations. If the initial A375 population were composed of a mixture of subpopulations with stably different migratory capacities, we would expect that each of these clonal populations would migrate at their own stable rate throughout a sequential migration experiment, either inherently poor, moderate, or proficient migrators. Instead, we found that all four clonal populations initially showed very low constricted migration rates, but over the sequential rounds of migration through 5 µm pores, all generated a Bottom-5 population that was more highly migratory (Figure S9A). Thus, even when a clonal population is isolated that is initially poorly migratory, a highly migratory population can be isolated by sequential rounds of constriction. We next asked whether the nucleus and 3D genome features of sequentially constricted cells were shared across these separate clones. Like the parental population, Bottom-5 cells from the clones showed a wrinkled, more variable Lamin A/C distribution after constriction (Figure S9B). We also observed some of the same compartment switches after constricted migration in the clones (Figure 5A and B). Different clones varied in the amount of 3D genome structure change they exhibited after constriction (Figure 5C), but all clones showed compartment shifts in the same direction for regions that switched compartments in A375-P (Figure 5D). Indeed, while the final chromosome conformations after constriction for the different clones were not all the same, we found that all the Bottom-5 compartment profiles clustered together, away from the initial and unconstricted Top-5 conditions for each clone (Figure 5E). Thus, after sequential constricted migration, cells shared more of their chromosome compartment structure features with constricted cells of other clones than they shared with the clonal population from which they were derived. Additionally, each clonal population experienced a weakening of B compartment interactions and often a gain of A compartment interactions after constriction, as observed in A375-P and MDA-MB-231 cells (Figure 5F). The t(5,13) translocation observed after constricted migration in A375-P did not occur in these clonal populations (Figure S9C and E), but we saw the emergence of a different translocation (t(2,5)) in clone 2 and the loss of a t(2,19) translocation in clones 3 and 4 after sequential constriction (Figure S9D and E). Two of the clones also exhibited copy number loss of chromosome 13 after constriction, as was observed in the parental A375 cells (Figure S9F).

**Figure 5.**
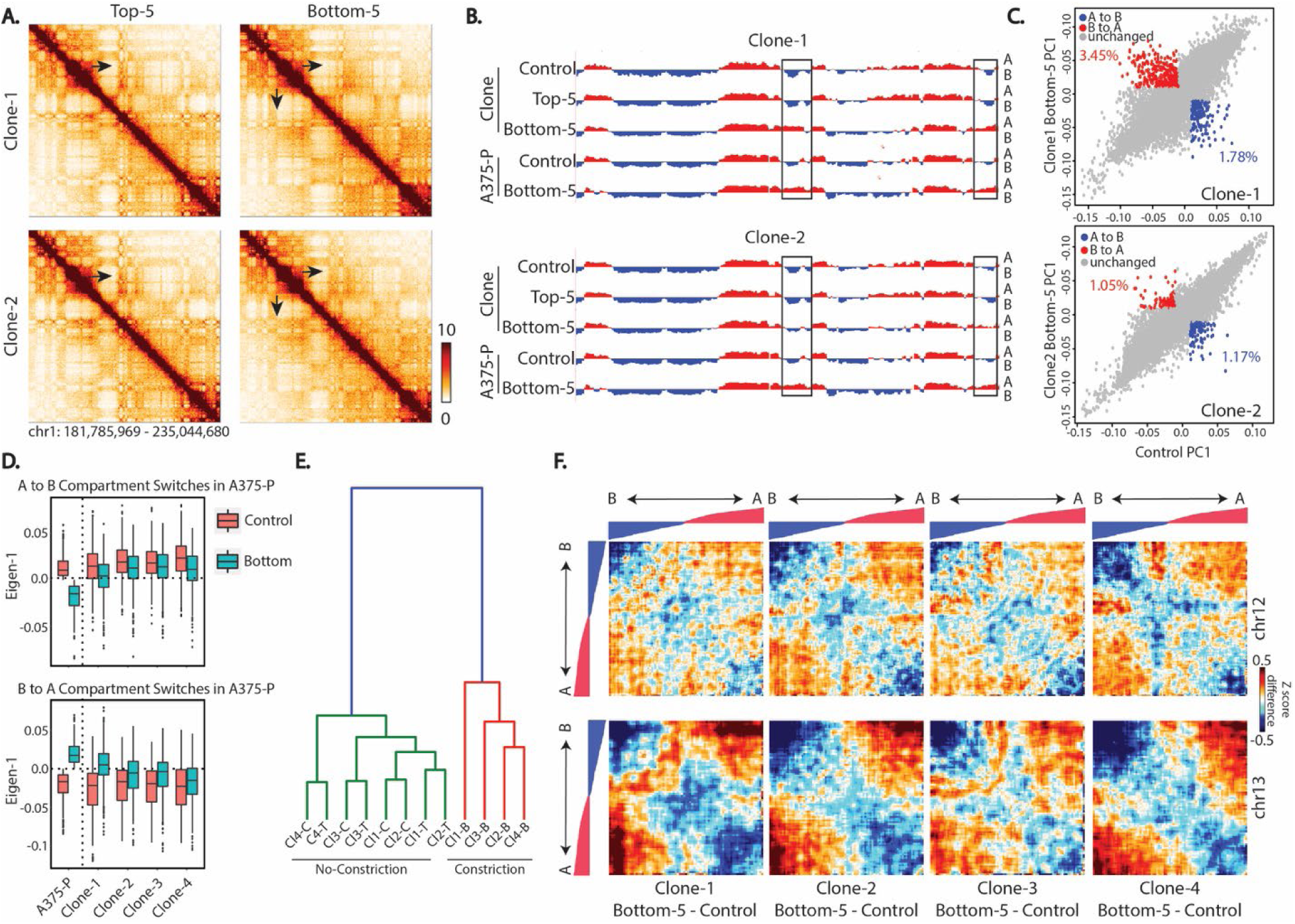
Compartment changes observed after constricted migration of A375 clonal populations. (A) 250 kb binned Hi-C interaction heatmaps of chr1 (chr1:181,785,969 - 235,044,680) in Clone-1 and Clone-2. Zoom-in region remains the same as in Figure 2A. Arrows: locations of visible difference in Bottom-5 contact maps for Clone-1. (B) PC1 track of compartment identity (250 kb bins) in the same region of chr1 across A375 clones and parental population (A375-P). Boxed region highlights the same compartment switch highlighted by arrows in A. (C) Compartment PC1 value comparisons for each bin genome wide in Clone-1 and Clone-2 Bottom-5 vs. Control. Percentage of bins that met the criteria for “compartment switch” from B to A (red; PC1 < -0.01 to > 0.01) or A to B (blue; PC1 > 0.01 to < -0.01) indicated on each panel. (D) “Eigen-1” = PC1 compartment values of each clone condition across regions that switch compartment identity in parental population from A to B (top graph) or B to A (bottom graph). (E) Clustering of all A375 clonal populations by their genome-wide compartment PC1. (F) Compartmentalization saddle plots of chr12 (top panels) and chr13 (bottom panels) display Z-score difference between all bin pairs of clonal Bottom-5 and clonal Control. Heatmaps are reordered by PC1 values from strongest B-B (left corner) to strongest A-A (right corner) as displayed by the overlaid tracks.

Genomic instability in cancer means that the clonal populations we examine will not stay entirely identical to the initial single cell, but by reducing the initial heterogeneity in the population, we observed common trends in 3D structure change that arise despite different starting points. While selection may act on heterogeneity that arises in each clonal population, it is not the case that constricted migration simply isolates stable different subtypes of cells from the A375-P population.

## Discussion

Our results identify different 3D genome structures specific to cells that have passed through numerous constrictions smaller than their nucleus. We find that such changes primarily occur at the A/B spatial compartmentalization level. A set of genomic regions show a switch in compartmentalization in both sequentially constricted A375 melanoma cells and MDA-MB-231 breast cancer cells. Beyond compartment switches, we observe a weakening of interactions in the typically heterochromatic B compartment that is consistent between both melanoma and breast cancer cell lines. These 3D genome structure differences in A375 cells are stable and associated with notable differences in cell and nucleus phenotypes both when the cells are migrating on a 2D surface and when they are squeezing through a 3D collagen matrix. Cells proficient at constricted migration more often deform their nuclei, elongate their cell body, and have fewer cell-cell attachments. These observations raise the interesting possibility that the increased migratory efficiency of sequentially constricted cells can be epigenetically encoded and stabilized by chromosome compartmentalization.

A key question raised by our results is whether the differences in 3D genome structure we see after constricted migration result from a selection process on an initially heterogeneous population or a “training process” where constricted migration itself leads to chromosome structure changes. Modeling results indicate that either selection, training, or a combination could explain the differences observed in sequentially constricted cells. The ability to isolate A375 cells that do not migrate in ten chances through a wide pore (Top-12 cells), shows that the initial A375 population contains some cells with a pre-existing migration phenotype that can be selected for by the sequential migration experiment. However, our clonal population results indicate that the A375 parental population is not simply composed of multiple subpopulations with stably different migratory potential. Each of the isolated clones start with low migration through constrictions, but a highly migratory population can be generated after rounds of sequential constricted migration. While these sequentially constricted clones do not all change their 3D genome in exactly the same way, they share common shifts in chromosome compartmentalization and a loss of the strongest B compartment interactions. These results indicate that the 3D genome structures characteristic of proficient constricted migrators are either selected and stabilized from a set of spontaneously arising fluctuations in chromosome structure or induced by the constricted migration. Similarly, some structures and phenotypes of Bottom-5 cells, such as the chromosome 5;13 translocation and extremely elongated cell shapes in A375 cells, and several gained translocations in MDA-MB-231 cells, are not detectable in the original control population and are thus likely the result of induced phenotype changes or random events that are further reinforced by a selection process. Overall, while our results cannot be explained by simple selection from a mixture of static pre-existing subpopulations, both selection and induced changes could contribute to the final phenotype. This combination of selection and induction likely occurs in metastasis as well. Within melanoma lesions, it has been observed that a select group of cells can gain the ability to metastasize, and then further diversification and phenotype change can occur as a result of metastasis (Damsky *et al*, 2010).

How could constricted migration induce chromosome conformation changes? One possible contributor could be physical deformation-induced pulling apart or pushing together of chromosomal regions. Mixing of chromosome regions during constriction might lead to new spatial associations due to coalescence of compartments in a phase separation-related process (Laghmach *et al*, 2021; Lee *et al*, 2021). Highly specific and reproducible local compartment switches, in contrast, are unlikely to have been pushed or pulled in the same way in every individual cell. Such specific changes could arise either by selection of pre-existing differences in the population, selection of a change that is induced by deformation in some cells during migration, or induction of a programmed cell response to constricted migration. It is known that mechanical signaling can induce cellular reprogramming and that cellular reprogramming, such as an epithelial to mesenchymal cell transition (Shivashankar, 2019), can involve genome structure changes. Constriction may also influence spatial compartments by altering chromatin modifications. Previous work has shown in other cell types that constricted migration can lead to an increase in heterochromatin (Hsia *et al*., 2021). We do not observe stable, long-term increases in heterochromatin marks in our sequentially constricted cells, but it is possible that more transient constriction-induced heterochromatin alterations could contribute to some of the stable compartment changes that we do observe.

We consistently observe loss of B-B interaction strength with constricted migration. Notably, this phenomenon was also previously observed with constricted migration in neutrophils (Irianto *et al*., 2017b). Further, we observe this loss of B-B interaction strength in the purely physical perturbation of low-salt expansion in isolated nuclei (Liu & Dekker, 2021; Sanders *et al*, 2022). Conversely, in lung cancer cells, the loss of SETDB1 methyltransferase was shown to leads to an increase in B-B compartment interactions and increased segregation of compartments overall, while also stiffening the nucleus and decreasing cell migration (Zakharova *et al*., 2021). This result is consistent with our findings in that we observe increased cell migration associated with the opposite changes in compartment interactions (loss of B-B compartment interactions and increased mixing of B-A). These coherent results across systems suggest a consistent relationship between chromosome compartment organization and nucleus mechanics and deformation.

Our results suggest that the constricted migration-specific 3D genome structure relates to several biological processes. Some constriction-specific spatial compartmentalization changes likely reinforce altered expression of migration-specific genes. While local gene regulation without 3D structure change can be sufficient to explain temporary changes in cellular behaviors in response to a stress or stimulus (Jin *et al*, 2013), 3D genome rearrangement is less reversible and more typical of differentiation or stable cell state transitions. Constricted migration has previously been shown to influence stem cell differentiation (Smith *et al*, 2019). Our results suggest that differentiation-like stable phenotype changes after constriction may be mediated by chromosome compartmentalization changes. Such stable differences in 3D genome structure after multiple constricted migration events could underlie the distinct phenotype and increased *in vitro* invasiveness of cancer cells derived from metastatic sites compared to those from primary tumors or normal tissue (Oppenheimer, 2006).

Some genomic regions with altered compartmentalization in sequentially constricted cells do not show a connection to gene regulation. In fact, we report an enrichment A to B compartment switches in regions with no genes. These changes could have a role beyond gene regulation, perhaps modulating the physical properties of the cancer cell nucleus. Previous research has shown that nucleus stiffness can be altered by increasing or decreasing the overall expression of Lamin A/C or by changing overall levels of heterochromatin (Davidson *et al*., 2014; Lammerding *et al*., 2006; Stephens *et al*., 2017; Stephens *et al*., 2018). However, our sequentially constricted cells showed no changes in overall Lamin A/C content or H3K9me3 heterochromatin levels even though their nuclei showed more deformations when embedded in a collagen matrix. Instead, we observed a redistribution of Lamin A/C and H3K9me3 localization in the nucleus. This suggests that altered distributions of these proteins and chromosome regions could contribute to the ability of the nucleus to undergo physical deformations. Future work will be required to determine whether the 3D genome changes we observe translate into quantitative differences in nucleus physical properties.

Our data also provide insights about basic 3D genome organization principles. We show that compartments, compartment strength, and whole chromosome interactions change while TADs remain unaltered, reinforcing the idea that these are separate and largely independent layers of genome organization (Mirny *et al*, 2019; Schwarzer *et al*, 2017). These changes also suggest that the compartment and whole chromosome level of genome structure may be more important to nuclear mechanics than the TAD and loop level of organization. Meanwhile, our observation that local punctate interactions change around sites where gene expression changes echo previous evidence that collisions between loop extrusion and transcription can influence local contact patterns (Brandao *et al*., 2019).

Could the 3D genome changes we observe in our constricted cell population be indicative of metastatic potential? Previous reports have sometimes detected gene expression signatures in primary tumors that may be predictive of metastasis (Ramaswamy *et al*, 2003). Nuclear morphology abnormalities (“nuclear atypia”) are also already used clinically as a marker of cancer aggressiveness (Kadota *et al*, 2012). Recent reports have taken advantage of microfluidic engineering to predict metastatic potential of breast cancer cells based on their morphological features (Yankaskas *et al*, 2019). But, beyond being a marker, the biological significance of such abnormalities and what chromosome structure changes accompany them have not been defined. It is possible that chromosome structure changes could link metastatic gene expression signatures with the abnormal nuclear appearance of aggressive cancer. Our results reveal chromosome spatial compartmentalization differences in genomic regions related to metastatic potential in sequentially constricted A375 cells. Future work will be needed to investigate whether related changes are also observed in cells from metastatic cancer in patients.

## Glossary

Control: Cells that have not undergone constricted migration but have been growing in a 2D culture dish.
Top-5: Cells that have not undergone constricted migration through 5 µm pores even after 10 chances to do so.
Bottom-5: Cells that have undergone constricted migration through 5μm pores for 10 rounds.
Top20-5: Cells that have not undergone constricted migration through 5 µm pores even after 20 chances to do so.
Bottom20-5: Cells that have undergone constricted migration through 5 µm pores for 20 rounds.
Top-12: Cells that have not undergone migration through 12 µm pores even after 10 chances to do so.
Bottom-12: Cells that have undergone migration through 12 µm pores for 10 rounds.

## Acknowledgements

We thank Rebeca San Martin, Mariano Labrador, Jacob Sanders, Yang Xu and Jan Lammerding for insightful discussion and advice about the project and methods. We thank Bas van Steensel for providing the LAD type annotations. Confocal images were obtained at the University of Tennessee Advanced Microscopy Imaging Core. This research was supported in part by a Ralph E. Powe Junior Faculty Enhancement Award from Oak Ridge Associated Universities to R.P.M. and by NIH NIGMS grant R35GM133557 to R.P.M. R.G. was supported in part by a Yates Dissertation Fellowship from UTK. D.T. was supported in part by a UTK Summer Undergraduate Research Internship award.

## Author Contributions

R.G. and R.P.M. conceived the project, designed the experiments, and wrote the paper with input from all authors. R.G. performed genomic, cell culture, and imaging experiments and bioinformatics data analysis. C.P. performed A375 clone Hi-C experiments, mFISH experiments, and some immunofluorescence experiments and analysis. T.F. performed the MDA-MB-231 sequential migration experiments and assisted with A375 experiments and Hi-C. P.D. designed and implemented the agent based modeling and assisted with computational analysis. T.I.R. performed 3D collagen immunostaining and quantification. J.H.G. performed translocation analyses. D.T provided experimental assistance and performed single cell migration video analysis.

---

**Supplementary Figure 1.**
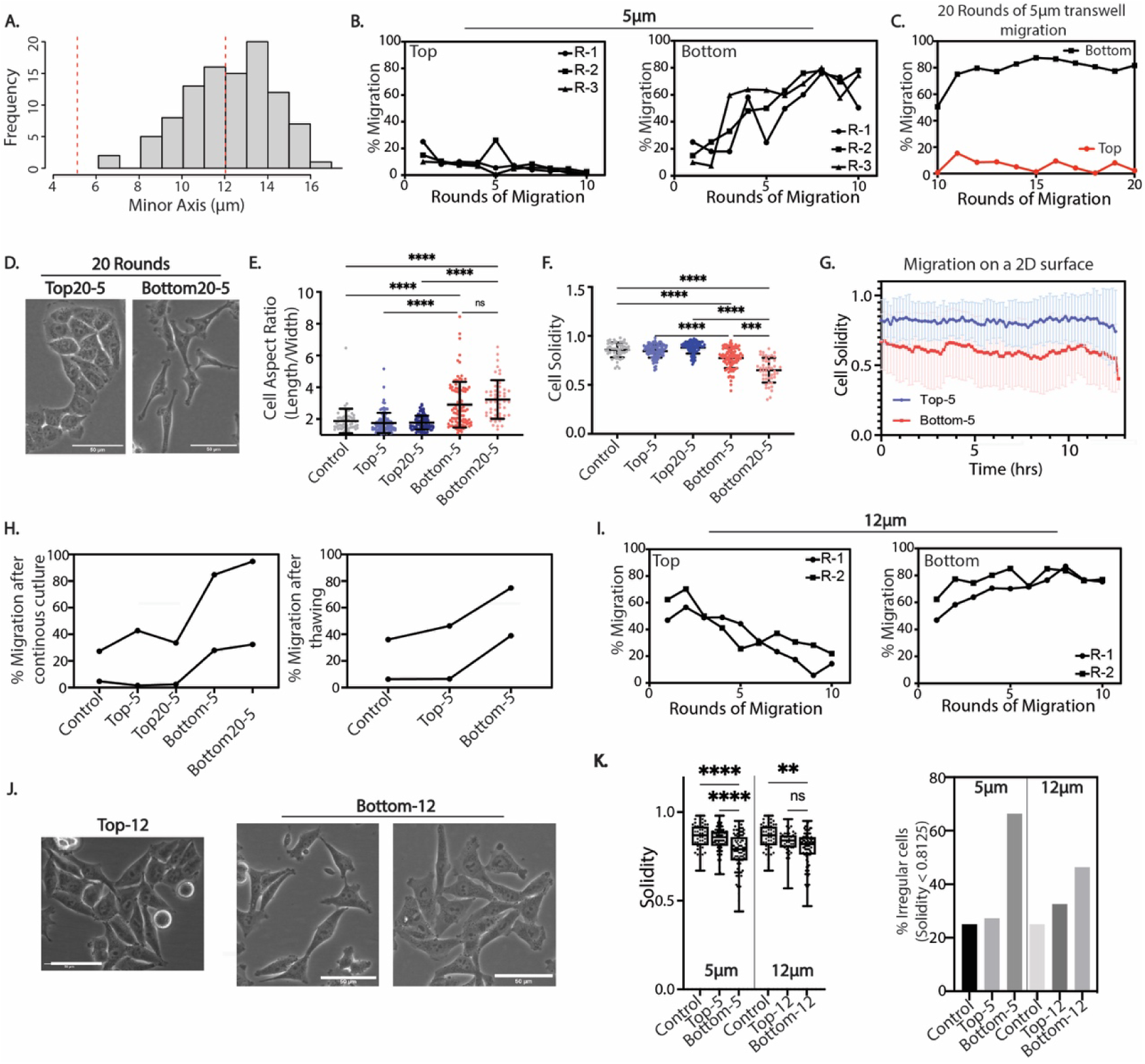
A375 cells exhibit persistent increase in migration efficiency and morphological changes after 5 µm constricted migration. (A) Minor axis (minimum diameter) measurements of A375 cell nuclei based on confocal z-stacks. N=100. Dotted lines indicate the sizes of pores (5 and 12 µm) used in the Transwell devices (B) Migration efficiency of each biological replicate for 10 rounds of 5 µm Transwell migration for Top (left) and Bottom (right) cells. (C) Migration efficiency of A375 cells through 20 rounds of 5 µm Transwell migration. Always Top (red) and Always Bottom (black) isolated in each round of migration as in B. (D) Phase contrast images of A375 cells subjected to 5 µm Transwell migration for 20 rounds. Scale = 50 µm. (E) Cell aspect ratio measurements of A375 cells that have undergone 10 rounds (Top-5 and Bottom-5) or 20 rounds (Top20-5 and Bottom20-5) of migration through 5 µm constrictions (**** p < 0.0001; Control (n = 53), Top-5 (n = 100), Top20-5 (n = 95), Bottom-5 (n = 100), Bottom20-5 (n = 51), Kruskal-Wallis multiple comparisons test) (F) Solidity measurements (proportion of object pixels in convex hull) of cells in all A375 subpopulations (**** p < 0.0001, *** p = 0.0003; Control (n = 52), Top-5 (n = 99), Top20-5 (n = 93), Bottom-5 (n = 98), Bottom20-5 (n = 48), Kruskal-Wallis multiple comparisons test) (G) Solidity measurements of A375 cells undergoing live-cell migration for 13 hours on a 2D surface (1 indicates ideally round cell shapes and lower values indicate more irregularities and protrusions). Error bars = mean ± SD for n = 14 Top-5 cells and n = 12 Bottom-5 cells (p < 0.0001, two-tailed t-test). (H) Migration efficiency of A375 cells after 5 passages of continuous culture (left graph) and after one cycle of freeze - thawing (right graph) in two biological replicates. (I) Migration efficiency of each replicate for 10 rounds of 12µm Transwell migration. (J) Phase contrast images of A375 cells after sequential 12 µm Transwell migration. Scale = 50 μm. (K) (Left panel) Cell solidity comparison after 5μm and 12μm pore migration. Box and whisker plot showing all data points in Control (n = 52), Top-5 (n = 99), Bottom-5 (n = 98), Top-12 (n = 43) and Bottom-12 (n = 82) (**** p < 0.0001, ** p = 0.0029, Kruskal-Wallis multiple comparisons test). (Right panel) % of A375 cells displaying irregular morphology (Solidity < 0.8125, first quartile of Control).

**Supplementary Figure 2.**
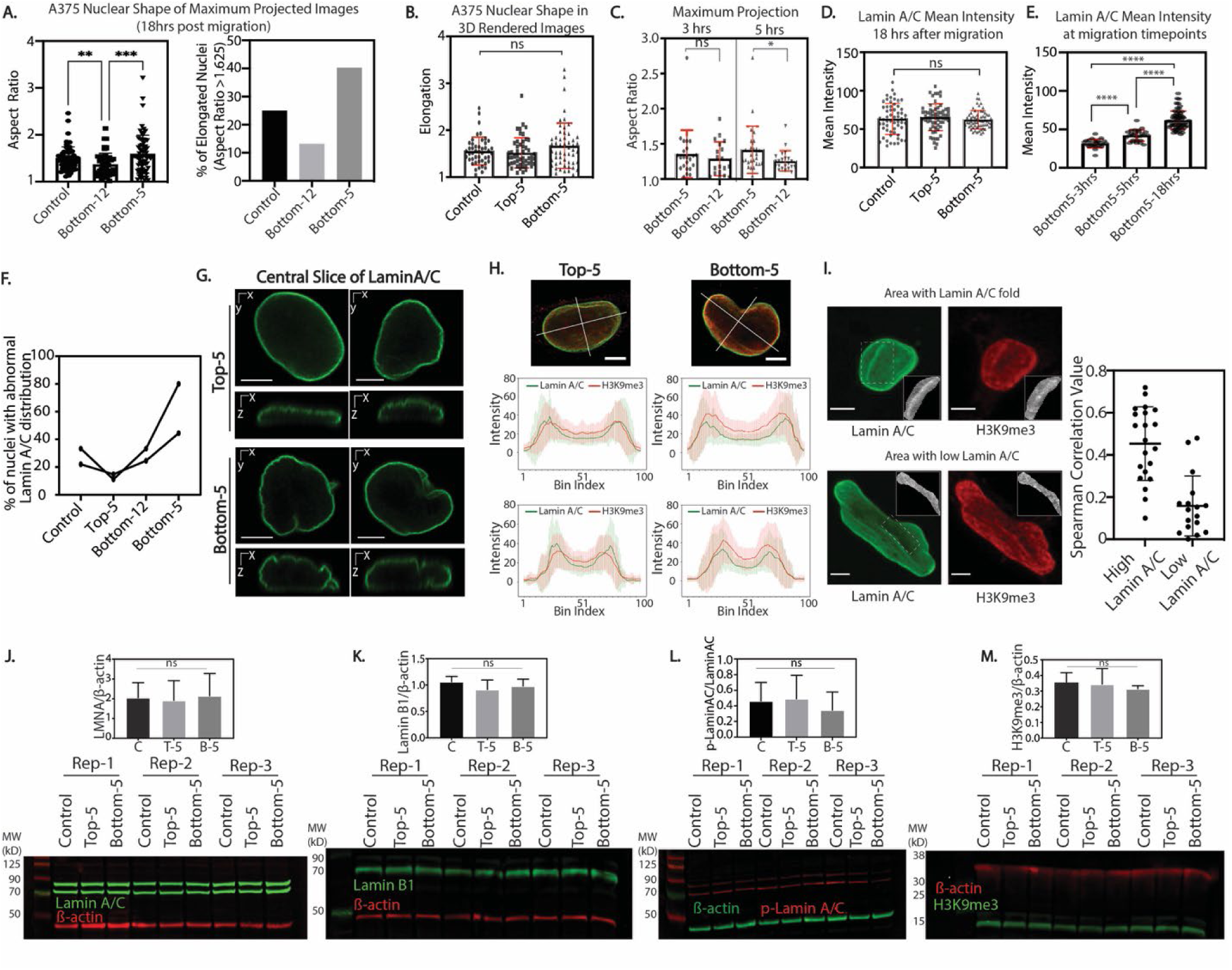
Changes in nuclear morphology after constricted migration associate with lamin and heterochromatin redistribution. (A) Left: Aspect ratio quantification of maximum projection images of A375 nuclei exposed to Transwell migration (** p =0.0099, *** p = 0.0004, Kruskal-Wallis multiple comparisons test; Control = 68, Bottom-12 = 53, Bottom-5 = 72 nuclei). Right: Fraction of A375 nuclei with Aspect Ratio >1.625 (3^rd^ quartile of Control). (B) Nuclear elongation quantification of 3D rendered A375 nuclei using NucleusJ in Control (n = 46), Top-5 (n = 50) and Bottom-5 (n = 54) cells. (C) Nucleus aspect ratio measurements at 3 and 5hrs after migrating through 5 µm Transwell pores (Bottom-5; n = 23 for 3hr and n = 26 for 5hr) and 12 μm (Bottom-12; n = 19 for 3hr and n = 22 for 5hr). (D) Lamin A/C mean intensity, quantified from maximum projection of immunofluorescence images, in A375 Control, Top-5, and Bottom-5 cells (Control = 59, Top-5 = 64, Bottom-5 = 62 nuclei). (E) Lamin A/C mean intensity in A375 cells 3, 5 and 18 hr after migrating through 5μm Transwell pores. Number of nuclei quantified remains the same as in C and D (**** p < 0.0001, two-tailed t-test). (F) Fraction of A375 cells displaying irregular patterns of Lamin A/C, defined by visual inspection of confocal images, in Control (n = 59), Top-5 (n = 63), Bottom-12 (n = 80) and Bottom-5 (n = 64) cells. Two biological replicates shown in the figure; lines connect A375 conditions within the same replicate. (G) Central orthogonal slices of Lamin A/C stained nuclei in Top-5 and Bottom-5 cells. (scalebar = 5 μm) (H) Intensity line scans across major and minor axes of Lamin A/C and H3K9me3 central slices in A375 Top-5 (n = 63) and Bottom-5 (n = 64) nuclei. Error bars = mean ± SD. (scalebar = 5 μm) (I) Correlation of pixel intensities between Lamin A/C and H3K9me3 in Bottom-5 cells. Left: Examples of Lamin A/C and H3K9me3 correlation with (top) or without (bottom) Lamin A/C enrichment. Areas used to calculate Spearman correlations between Lamin A/C and H3K9me3 intensity indicated with dotted yellow outline and in grayscale inset. Right: Spearman correlation values between Lamin A/C and H3K9me3 across 20 nuclear areas with high Lamin A/C intensity (stripes) and 18 nuclear areas low in Lamin A/C. (J-M) Western blot analysis of A375-Control, Top-5 and Bottom-5 cells probing for Lamin A/C (J), Lamin B1 (K), phospho-Lamin A/C (Ser22) (L) and H3K9me3 (M). Quantification shown above each blot. No significant differences found in pairwise comparisons by two-tailed t-test.

**Supplementary Figure 3.**
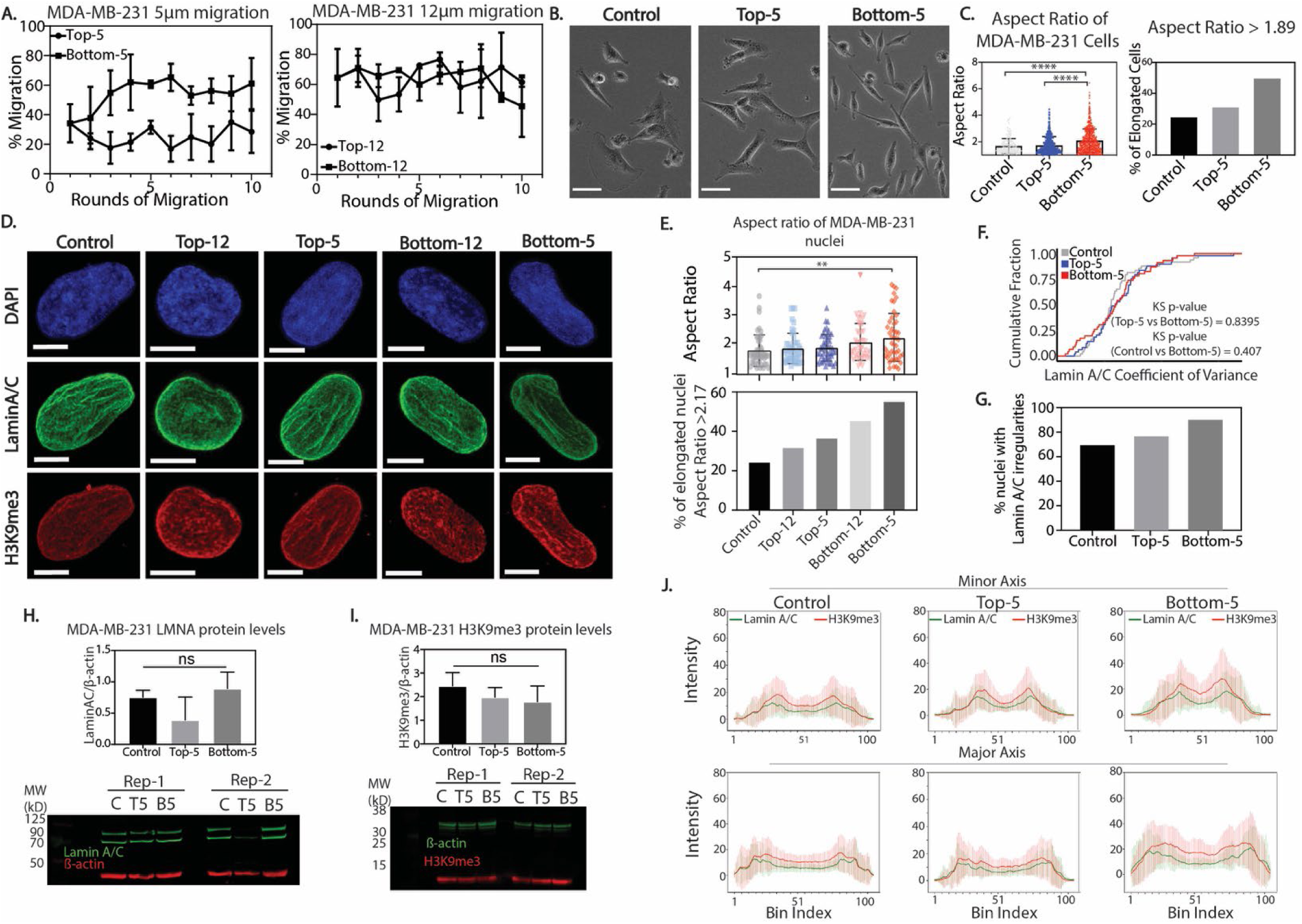
Sequential rounds of constricted migration in MDA-MB-231 cells. (A) Transwell migration efficiency of MDA-MB-231 cells through 5 and 12 µm pore sizes for 10 Transwell migration rounds. Error bars = mean ± SD of 5 μm (3 biological replicates) and 12 μm (2 biological replicates). (B) Phase contrast images of MDA-MB-231 cell subpopulations. Scale bars = 50 μm. (C) Left: Aspect ratio of MDA-MB-231 cells for Control (n = 142), Top-5 (n = 393) and Bottom-5 (n = 621) (**** p < 0.0001; two-tailed t-test). Right: Fraction of MDA-MB-231 cell subpopulations with an aspect ratio > 1.89 (3^rd^ quartile of Control). (D) Maximum projected images of MDA-MB-231 nuclei immunostained with DAPI (blue), Lamin A/C (green) and H3K9me3 (red). Scale bars = 5μm. (E) Top: Aspect ratio of MDA-MB-231 nuclei in Control (n = 49), Top-12 (n = 47), Top-5 (n = 49), Bottom-12 (n = 46) and Bottom-5 (n = 47) (** p = 0.0038; two-tailed t-test). Bottom: Fraction of MDA-MB-231 nuclei that have an aspect ratio > 2.17 (3^rd^ quartile of Control). (F) Cumulative distribution plot of Lamin A/C coefficient of variance in Control, Top-5 and Bottom-5 MDA-MB-231 cells. Number of nuclei quantified remains the same as in E. KS-test shows no significant difference between conditions. (G) Fraction of MDA-MB-231 nuclei displaying irregularities in Lamin A/C distribution by visual inspection of confocal images. (H-I) Western blot analysis and quantification of MDA-MB-231 Control, Top-5 and Bottom-5 cells probing for Lamin A/C and H3K9me3. Error bars = mean ± SD. No significant differences found in pairwise comparisons by two-tailed t-test. (J) Line scan intensity for H3K9me3 (red) and Lamin A/C (green) across major and minor axis of central slice images of MDA-MB-231 cells. Number of nuclei quantified remains the same as in E. Error bars = mean ± SD.

**Supplementary Figure 4.**
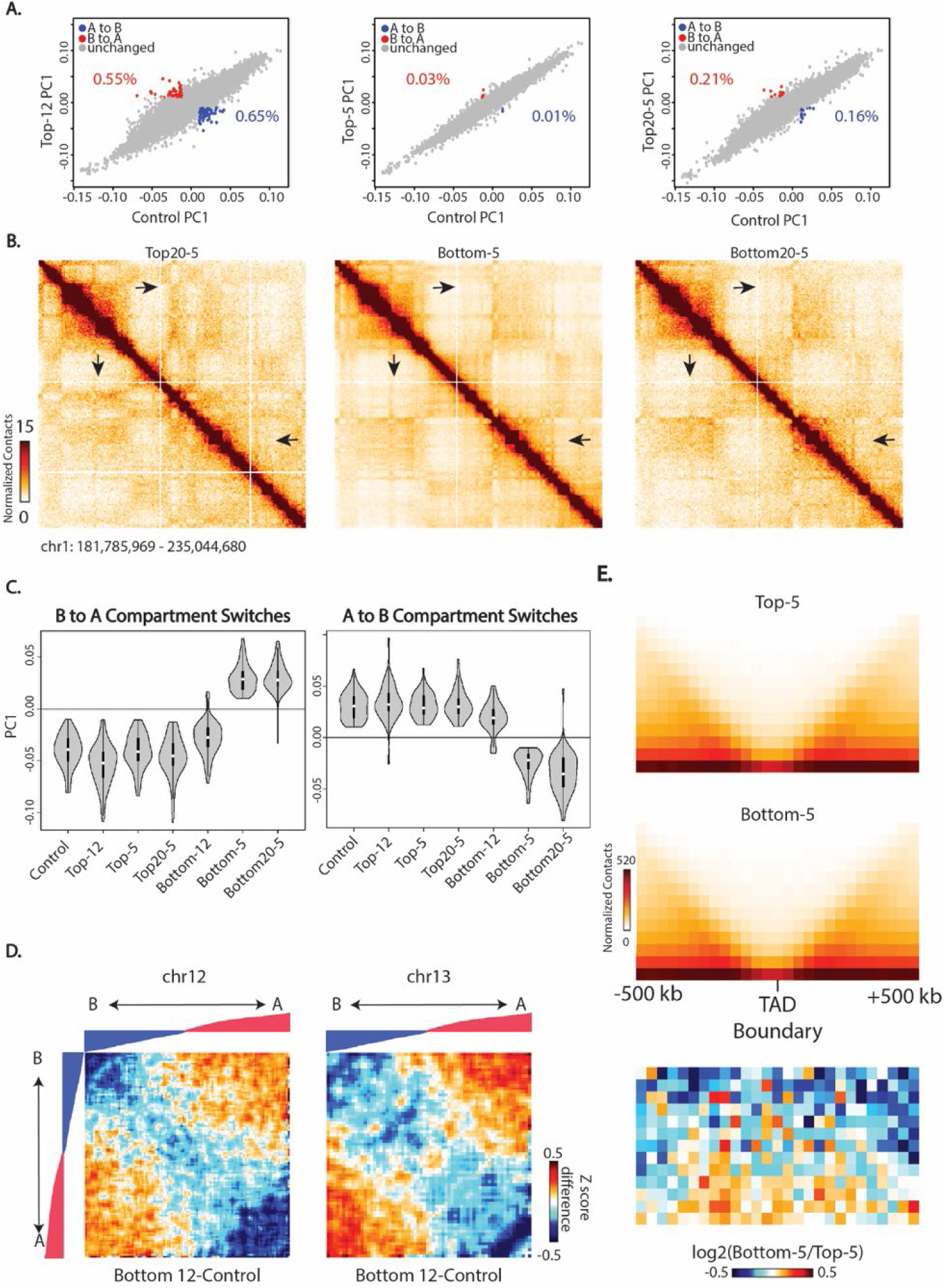
Compartment but not TAD changes are consistently associated with constricted migration. (A) Compartment PC1 value comparisons for each bin genome wide for cells that did not migrate through 5 or 12 µm pores. Significant compartment switches indicated in red or blue with associated percentages of bins. (B) Hi-C interaction heatmaps of chr1 (same chromosomal location as Figure 2A) in A375 cells that have undergone 20 rounds of 5 µm Transwell migration. Black arrows indicate regions that exhibit changes in compartment profiles upon migration through 5 µm constrictions. (C) Distribution of PC1 values across each A375 subpopulation for genomic regions that switched their compartment identity upon 5 µm constricted migration from B to A compartment (left panel) or A to B compartment (right panel). (D) Change in interaction Z-score from Control to Bottom-12 for chr12 (left) and chr13 (right). Bins are reordered based on A375 Control PC1 values also represented as tracks above the heatmaps. (E) Aggregated 40 kb binned contact maps at TAD boundaries called by the insulationScore method with strength greater than 1. Bottom panel: log2ratio of the Bottom-5 and Top-5 aggregated contacts around the same TAD boundaries.

**Supplementary Figure 5.**
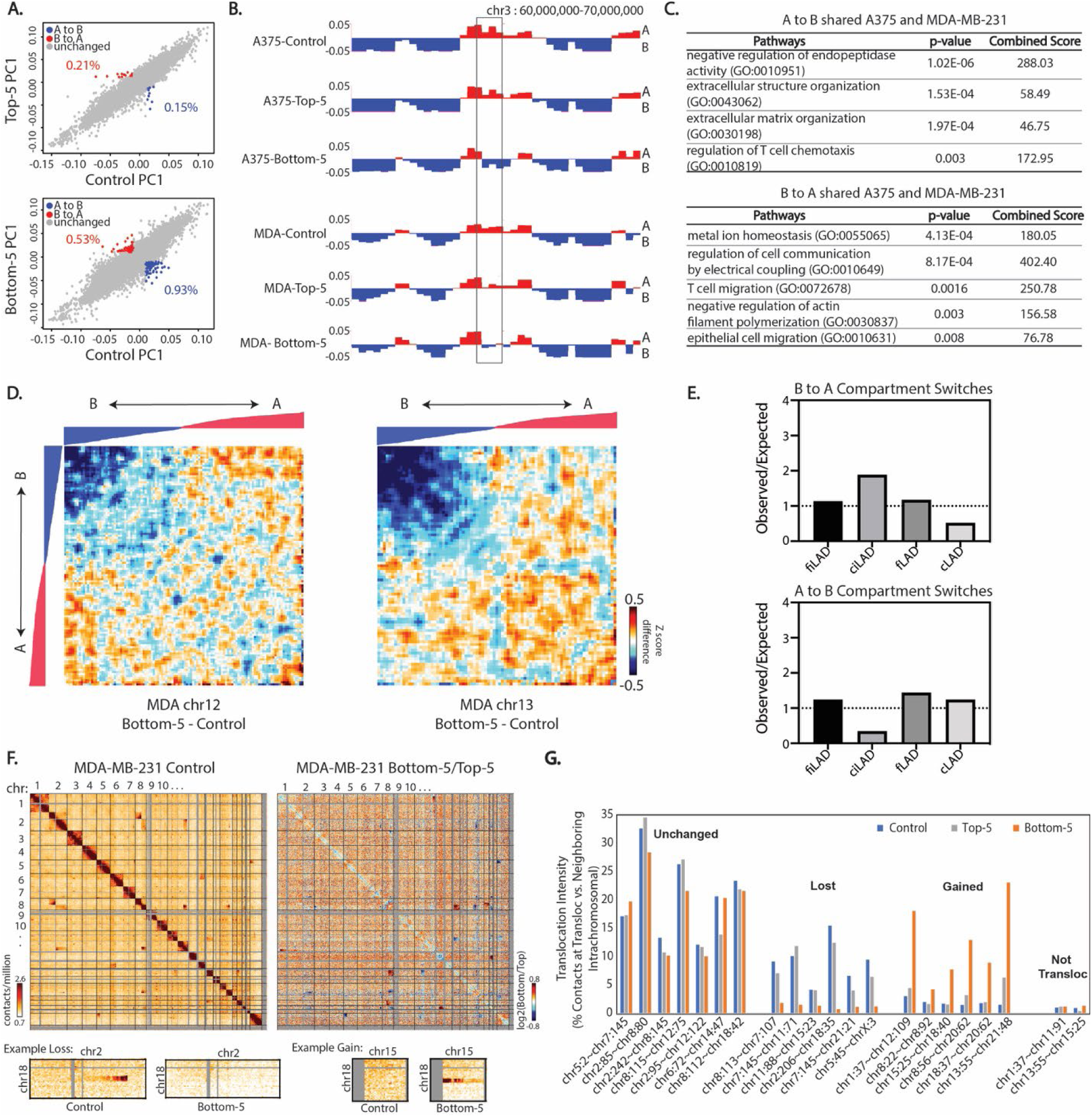
Constricted migration is associated with genome structure differences in MDA-MB-231 cells. (A) Scatterplots of PC1 compartment eigenvector values show compartment changes (red = B to A and blue = A to B) between Control MDA-MB-231 cells and either Top or Bottom cells from 10 rounds of 5 µm Transwell migration. (B) PC1 compartment track (250 kb bins) in A375 and MDA-MB-231 cells. Box highlights example region that switches from A to B compartment in both A375 and MDA-MB-231 cells in 5 µm constricted migration. (C) Pathway analysis of genes found within the compartment switch regions shared between A375 and MDA-MB-231. (D) Compartmentalization saddle plots of chr12 (left) and chr13 (right) display interaction Z score differences between Bottom-5 vs. Control MDA-MB-231 cells. 250 kb bins, excluding those that show compartment changes, are sorted by MDA-MB-231 Control PC1 values. (E) Enrichment of different LAD types in regions that switch compartments in Bottom-5 MDA-MB-231 cells compared to Control. (F) Whole genome 2.5 Mb Hi-C interaction frequency maps for MDA-MB-231 Control (left) and log2 ratio of MDA-MB-231 Bottom-5/Top-5. Lower panel heatmaps display zoom-in of regions with altered translocation signals. (G) Translocation intensities for translocations maintained, lost after constricted migration, or gained after constricted migration. “Not transloc” shows the calculated intensity for a control non-translocated interchromosomal region. See Methods for calculation details.

**Supplementary Figure 6.**
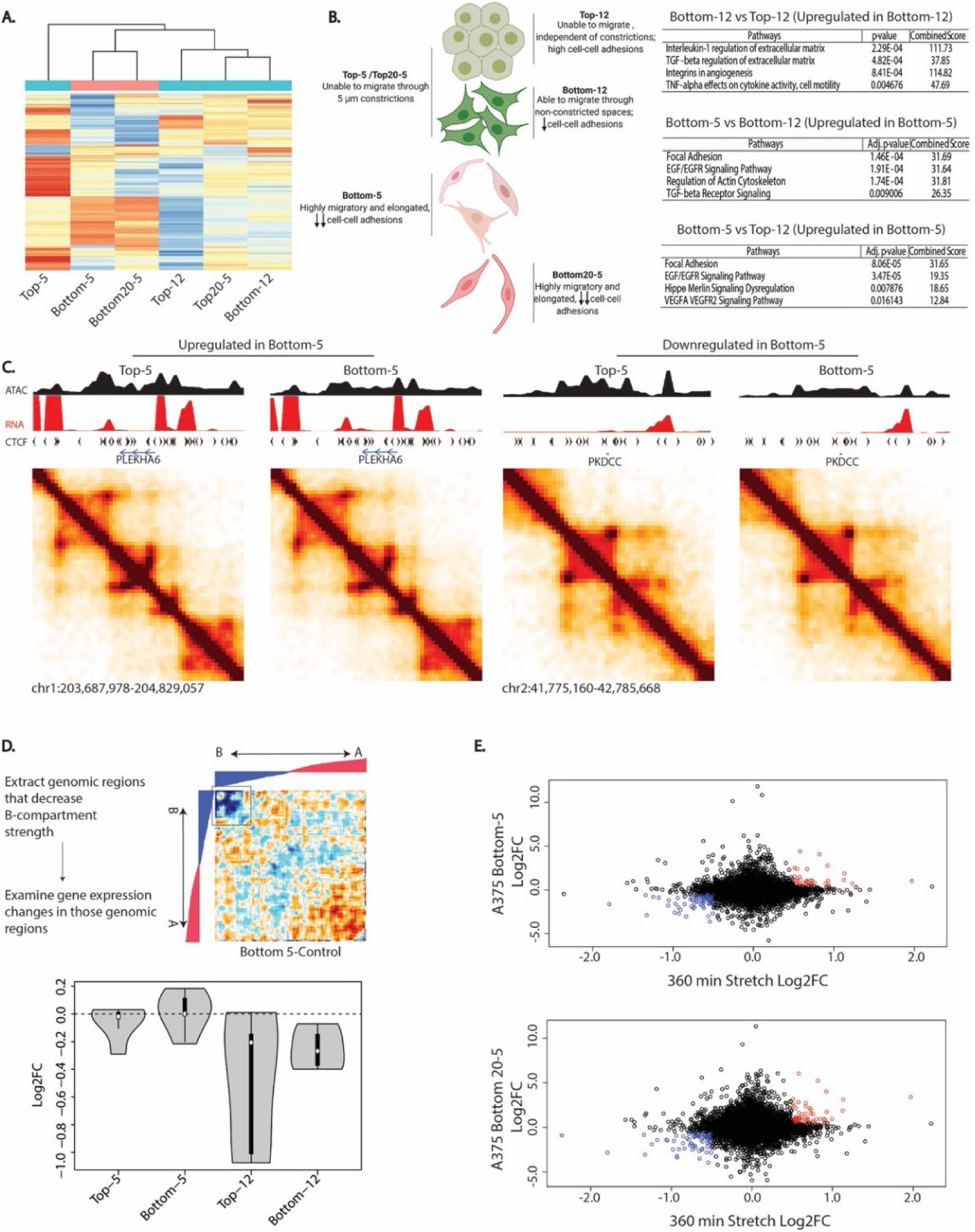
Distinct gene expression profiles of sequentially constricted cells. (A) Hierarchical clustering of all A375 subset of cells based on genes that are differentially expressed in Bottom-5 cells relative to Control (Log2FC >1). (B) Gene ontology analysis of differentially expressed genes comparing indicated sub-populations of A375 cells. Cell illustration indicates morphology differences we observe within each sub-population of A375 cells. (C) Hi-C contact matrices binned at 20 kb and smoothed at 40 kb bin size of upregulated genes (PLEKHA6, left two panels) and downregulated genes (PKDCC, right two panels) in Bottom-5 cells. Black track represents ATAC-Seq signal while the red track represents RNA-Seq signal. Black arrows indicate known CTCF motif directionality. (D) Genes corresponding to regions experiencing strong B compartment interaction loss in Bottom-5 cells genome-wide were analyzed for their changes in expression in indicated conditions vs. Control. (E) Correlation plots of differentially expressed genes between mechanically stretched cells (Nava *et al*., 2020) and A375 cells that have undergone constricted migration (10 rounds of 5 µm constriction, top; 20 rounds of 5 µm constriction, bottom). Red and blue dots indicate the few genes that change in the same direction in both conditions.

**Supplementary Figure 7.**
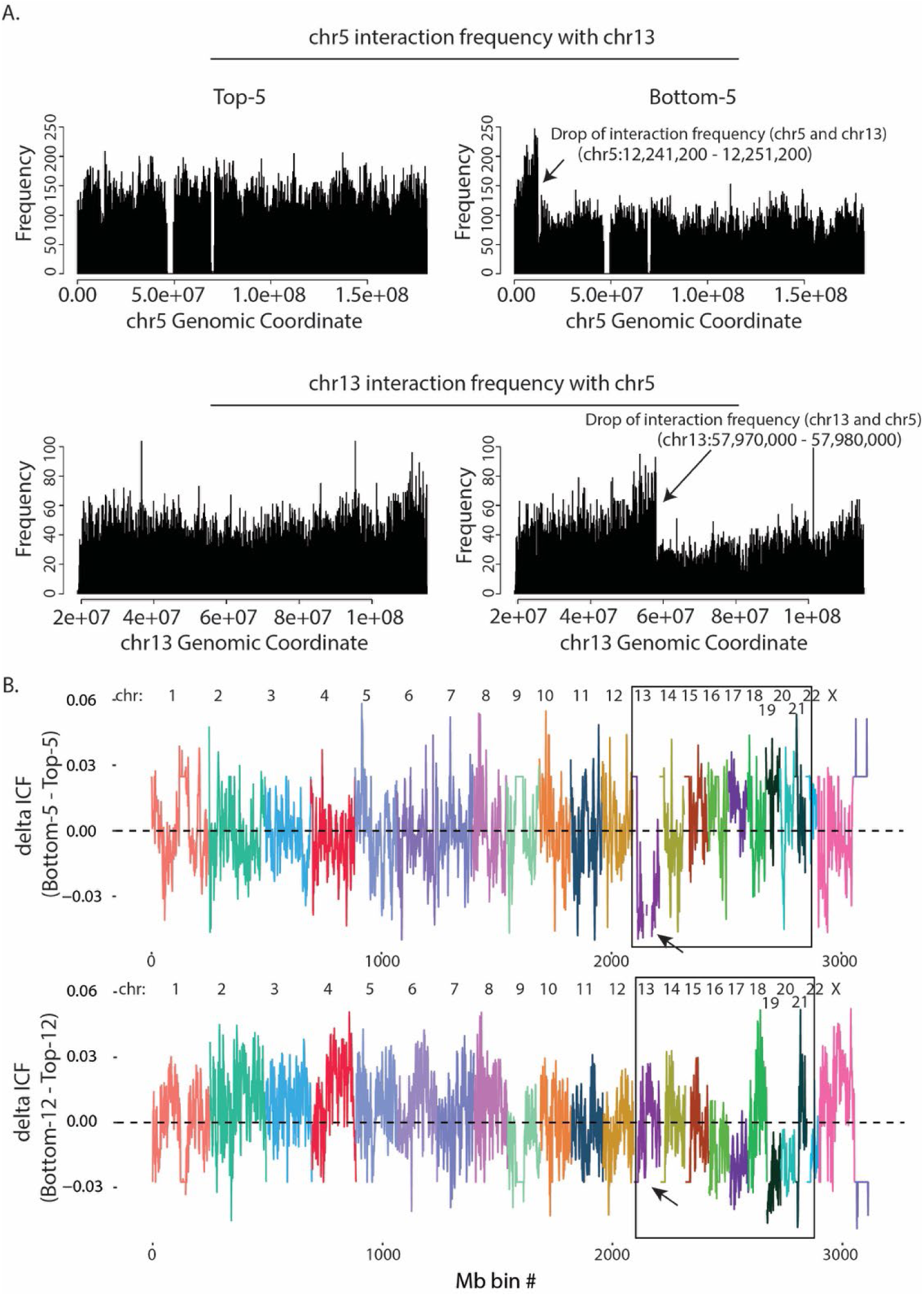
Identifying the breakpoint of the translocation between chr5 and chr13. (A) Top: Contact frequency of chr5 genomic coordinates with chr13 in Top-5 cells (left) and Bottom-5 cells (right). Bottom: Contact frequency of chr13 genomic coordinates with chr5 in Top-5 cells (left) and Bottom-5 cells (right). Black arrows indicate sudden drop in contact frequency indicating the site of the translocation. (B) Plots represent the difference in inter-chromosomal fraction (delta ICF) in cells that have undergone migration through 5 μm constrictions (top plot) and cells that have undergone migration through 12 μm constrictions (bottom plot) using 1Mb binned Hi-C contacts. The boxed region highlights regions of difference between constricted and unconstricted migration.

**Supplementary Figure 8.**
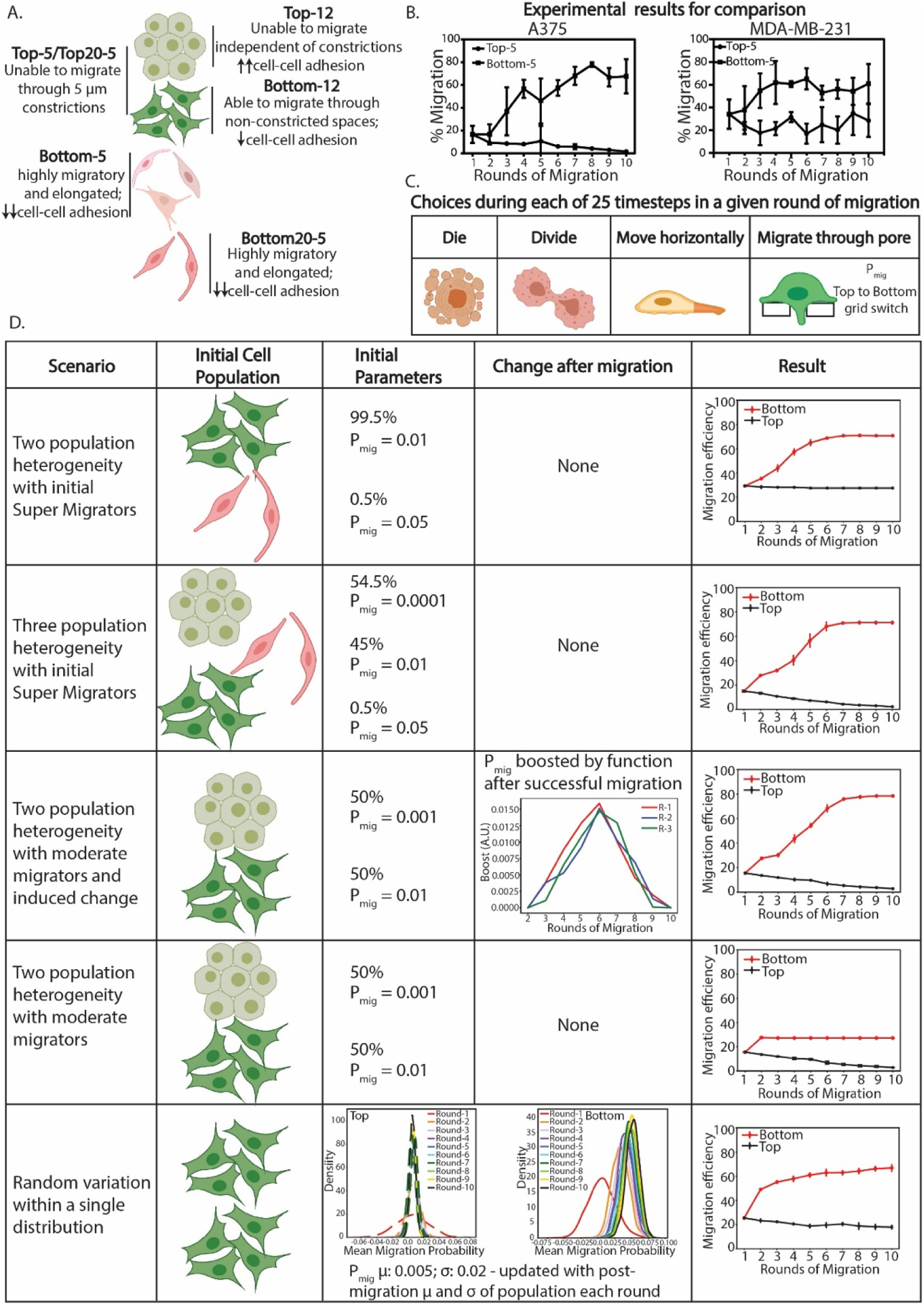
Agent based modeling of cell populations undergoing constricted migration. (A) A375 cell subpopulations observed after sequential constricted migration. (B) For model result comparison, results of sequential Transwell migration through 5μm constrictions are recapitulated from Figure 1 for A375 (left) and from Figure S3 for MDA-MB-231 (right) cells. (C) Choices a cell can make at each timestep during each simulated migration round. (D) Parameters used in each agent-based model (left columns) and results of each model (rightmost column).

**Supplementary Figure 9.**
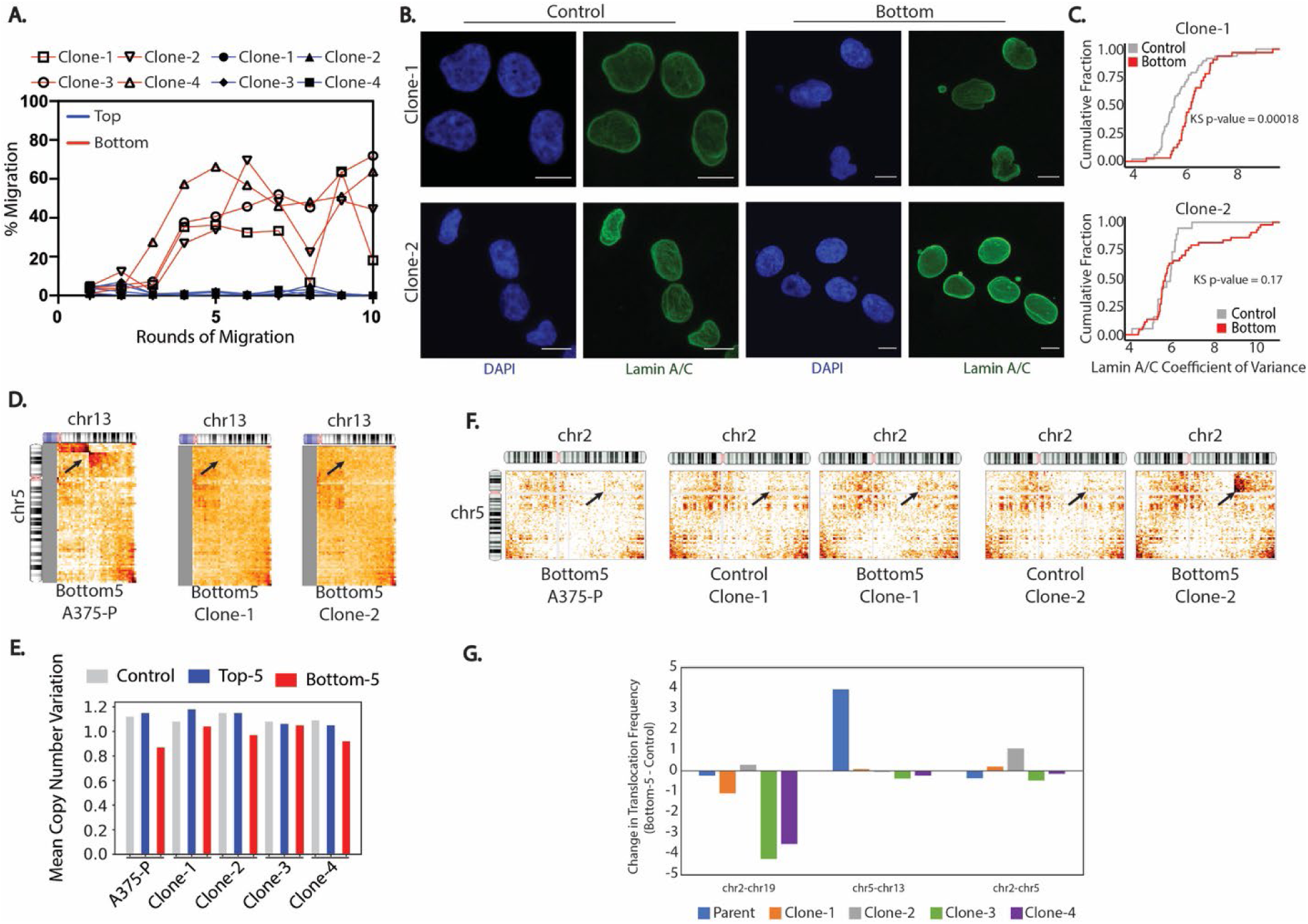
Sequential migration of A375 clones. A) Increase in migration efficiency is seen after sequential rounds of migration through 5 µm Transwell pores in all A375 clones. Cells that passed through pores each round = Bottom; red. Cells that did not migrate through pores = Top; blue. B) Sample images of Clone1 and Clone 2 A375 nuclei before (Control) or after (Bottom) 10 rounds of constricted migration. Nuclei stained with DAPI (blue) or Lamin A/C (green). Images represent a maximum projection of z-stacks (Scale bar= 5 µm). C) Cumulative distribution plot of Lamin A/C coefficient of variance. Kolmogorov-Smirnov test p-values for Bottom vs. Control Clone 1 (p = 0.00018, n = 48 nuclei for Control and 32 nuclei for Bottom-5) and Clone 2 (p = 0.17, n = 19 nuclei for Control and 44 nuclei for Bottom-5). D) Inter-chromosomal interactions between chr5 and chr13 (2.5 Mb bins) for parental population and clone cells that have undergone 5 μm Transwell migration. Arrow indicates site of translocation observed in parental population. E) Copy number variation of chr13 inferred from Hi-C data. The mean copy number across all samples is set at 1. Relative copy number loss is seen after constricted migration (Bottom-5 cells) for parental population as well as Clone 2 and 4. F) Inter-chromosomal interactions between chr5 and chr2 (2.5 Mb bins) for parental population and pre- and post-migration Clone1 and 2 cells. Arrow indicates site of translocation observed only in Bottom-5 Clone 2 cells. G) Translocation intensity for 3 different translocations (t(2;19), t(5;13), and t(2;5)) was quantified as contacts at the translocation site divided by average diagonal contacts for each Clone in Bottom-5 and Control. The change in this translocation intensity after constricted migration is plotted. Negative values indicate loss of the translocation after sequential constricted migration while positive values indicate translocation gain between the indicated chromosomes.

## Supplementary Tables

**Table S1.** Summary Statistics for Hi-C Sequencing Experiments

| Sample | Total Raw Reads | Both Sides Mapped | Valid Pairs | Unique Valid Pairs | % Cis | % Dangling Ends |
| --- | --- | --- | --- | --- | --- | --- |
| A375-Control-R1 | 201,352,157 | 124,670,760 | 123,653,686 | 107,358,325 | 61.2 | 0.35 |
| A375-Control-R2 | 200,881,087 | 102,710,892 | 101,397,252 | 80,394,938 | 47.7 | 0.31 |
| A375-Top10-5μ-R1 | 320,786,521 | 163,004,931 | 161,788,811 | 132,650,834 | 55.6 | 0.23 |
| A375-Bottom10-5μ-R1 | 386,733,576 | 193,207,211 | 191,853,175 | 152,440,505 | 60.3 | 0.21 |
| A375-Top10-5μ-R2 | 164,616,353 | 84,226,628 | 83,087,615 | 68,000,595 | 48.7 | 0.28 |
| A375-Bottom10-5μ-R2 | 107,320,517 | 54,380,776 | 53,662,580 | 43,911,557 | 49.2 | 0.45 |
| A375-Top20-5μ-R1 | 142,531,635 | 66,090,879 | 62,567,577 | 49,317,732 | 47.7 | 0.60 |
| A375-Bottom20-5μ-R1 | 159,577,275 | 73,992,021 | 70,695,806 | 57,329,882 | 49.7 | 0.44 |
| A375-Top10-12μ-R1 | 117,695,787 | 90,932,905 | 90,442,670 | 81,815,020 | 43.0 | 0.39 |
| A375-Bottom10-12μ-R1 | 110,637,334 | 85,346,366 | 84,975,979 | 76,728,534 | 37.0 | 0.30 |
| A375-Top10-12μ-R2 | 145,063,083 | 110,772,496 | 110,380,318 | 98,396,558 | 40.4 | 0.26 |
| A375-Bottom10-12μ-R2 | 126,055,079 | 96,662,104 | 96,161,324 | 86,277,860 | 43.5 | 0.34 |
| MDA-Control | 145,250,671 | 75,273,429 | 74,220,998 | 60,781,296 | 36.2 | 0.47 |
| MDA- Bottom10-5μ | 186,575,335 | 95,844,081 | 94,485,413 | 74,425,665 | 46.17 | 0.36 |
| MDA-Top10-5μ | 172,743,231 | 88,691,523 | 87,219,726 | 70,082,743 | 50.99 | 0.41 |
| A375-Clone1-Control | 415,896,902 | 218,953,001 | 205,096,854 | 146,952,851 | 81.3 | 4.06 |
| A375-Clone2-Control | 295,473,235 | 158,629,154 | 141,899,367 | 115,404,954 | 80.5 | 7.38 |
| A375-Clone3-Control | 118,321,345 | 68,964,948 | 57,710,790 | 47,997,040 | 75.6 | 12.51 |
| A375-Clone4-Control | 377,224,508 | 118,178,074 | 104,536,528 | 88,965,611 | 75.7 | 8.57 |
| A375-Clone1-Top10-5μ | 177,991,674 | 106,369,261 | 102,258,363 | 82,510,023 | 79.7 | 1.98 |
| A375-Clone2-Top10-5μ | 188,990,385 | 107,144,596 | 93,062,887 | 76,482,056 | 78.7 | 8.36 |
| A375-Clone3-Top10-5μ | 331,760,181 | 115,878,108 | 88,618,488 | 75,896,047 | 78.1 | 16.33 |
| A375-Clone4-Top10-5μ | 207,352,979 | 68,876,258 | 62,052,625 | 52,457,439 | 77.2 | 6.74 |
| A375-Clone1-Bottom10-5μ | 176,529,569 | 99,847,197 | 96,198,965 | 78,283,396 | 77.1 | 2.39 |
| A375-Clone2-Bottom10-5μ | 199,764,485 | 111,728,932 | 100,515,682 | 80,947,058 | 76.6 | 7.44 |
| A375-Clone3-Bottom10-5μ | 96,864,809 | 55,764,250 | 47,993,860 | 40,183,003 | 77.1 | 10.11 |
| A375-Clone4-Bottom10-5μ | 229,564,420 | 72,750,874 | 64,028,035 | 53,537,618 | 75.5 | 8.98 |

**Table S2.** Summary Statistics for RNA Sequencing Experiments

| Sample | Total Reads after rRNA removal | Mapped Reads | Percent Mapped Reads |
| --- | --- | --- | --- |
| Control-R1 | 25,802,932 | 23,454,526 | 90.9 |
| Control-R2 | 44,936,964 | 39,168,478 | 87.16 |
| Top5-R1 | 57,420,692 | 49,189,316 | 85.66 |
| Top5-R2 | 40,240,820 | 34,943,927 | 86.84 |
| Bottom5-R1 | 44,784,816 | 37,982,597 | 84.81 |
| Bottom5-R2 | 42,149,066 | 36,445,763 | 86.47 |
| Top20-5-R1 | 39,235,465 | 36,102,375 | 92.01 |
| Top20-5-R2 | 97,958,087 | 89,841,371 | 91.71 |
| Bottom20-5-R1 | 107,282,198 | 98,413,826 | 91.73 |
| Bottom20-5-R2 | 54,332,623 | 49,381,035 | 90.89 |
| Top12-R1 | 47,231,386 | 42,228,487 | 89.41 |
| Top12-R2 | 44,863,196 | 41,601,169 | 92.73 |
| Bottom12-R1 | 43,203,103 | 38,744,667 | 89.68 |
| Bottom12-R2 | 8,436,059 | 7,655,956 | 90.75 |

**Table S3.** Summary Statistics for ATAC Sequencing Experiments

| Sample | Total Reads | Reads after trimming | Mapped Reads | Percent Mapped Reads |
| --- | --- | --- | --- | --- |
| Control-R1 | 169,579,026 | 169,348,735 | 137,428,769 | 81.15 |
| Top5-R1 | 169,763,417 | 113,906,483 | 94,322,265 | 82.81 |
| Bottom5-R1 | 213,830,754 | 115,332,896 | 108,423,723 | 94.01 |

## Supplementary Movie Captions

**Supplementary Movie 1.** Phase contrast live cell imaging in a 2D culture dish of Top-5 cells in 3 different experiments. New frame every 10 minutes, total length = 13 hours.

**Supplementary Movie 2.** Phase contrast live cell imaging in a 2D culture dish of Bottom-5 cells from 3 different experiments. New frame every 10 minutes, total length = 13 hours.

**Supplementary Movie 3.** 3D rendering of Top-5 nucleus using Leica Sp8 software displaying Lamin A/C (green) and H3K9me3 (red). Scale bars (white) indicate a length of 2 μm.

**Supplementary Movie 4.** 3D rendering of Bottom-5 nucleus using Leica Sp8 software displaying Lamin A/C (green) and H3K9me3 (red). Scale bars (white) indicate a length of 2 μm.

**Supplementary Movie 5.** Live time-lapse imaging of a Top-5 cell embedded in 3D collagen matrix. Nuclei labelled with Dendra2-Histone H4.

**Supplementary Movie 6.** Live time-lapse imaging of Bottom-5 cells migrating through 3D collagen matrices. Nuclei labelled with Dendra2-Histone H4.

**Supplementary Movie 7.** 3D rendering of nucleus of collagen embedded Top-5 cell stained with DAPI (blue) displays a uniform distribution of Lamin A/C (green). Scale bars (white) indicate a length of 2μm.

**Supplementary Movie 8.** 3D rendering of nucleus of collagen embedded Bottom-5 cell stained with DAPI (blue) displays an elongated shape and an altered distribution of Lamin A/C (green) with appearance of fiber like structures on the side of the constriction. Scale bars (white) indicate a length of 5μm.

## MATERIALS and METHODS

## KEY RESOURCES TABLE

| REAGENT or RESOURCE | SOURCE | IDENTIFIER |
| --- | --- | --- |
| ***Antibodies (IF)*** |  |  |
| Lamin A/C | Santa Cruz | sc-376248 |
| Lamin B1 | Abcam | ab133741 |
| H3K9me3 | Abcam | ab176916 |
| Alexa Fluor 594 | Invitrogen | R37117 |
| Alexa Fluor 488 | Invitrogen | R37120 |
| ***Antibodies (Western Blot)*** |  |  |
| Beta Actin – Rabbit Polyclonal | Thermo Fisher | PA1-16889 |
| Beta Actin – Mouse Monoclonal | Thermo Fisher | MA1-140 |
| phospho LaminA Ser22 | Abcam | ab138450 |
| Goat anti-rabbit secondary | Licor | 92568071 |
| Goat anti-mouse secondary | Licor | 95-32210 |
| ***Chemicals*** |  |  |
| DMEM | Corning | 10-013-CV |
| FBS | Corning | 35-010-CV |
| Pen-Strep | Gibco | 14140-122 |
| L-Glutamine | Gibco | 25030-081 |
| Medium 199 | Lonza | 12-119F |
| Hygromycin B | Corning | 30-240-CR |
| Trypsin | Corning | 25053CI |
| RIPA buffer | Thermo Scientific | I89900 |
| Collagen | Corning | 354236 |
| protease inhibitors/EDTA | GenDepot | 50-101-5485 |
| phosphatase inhibitors | GenDepot | 50-101-5488 |
| micrococcal nuclease | Thermo Scientific | FEREN0181 |
| BCA Protein Assay Kit | Thermo | 234225 |
| 4-12% Bis-Tris Plus gels | Invitrogen | NW04120BOX |
| PVDF membranes | Invitrogen | B1000 |
| Chr5 Green Paint | Metasystems | D-0305-050-FI |
| Chr13 Orange Paint | Metasystems | D-0313-050-OR |
| ***Critical Commercial Assays*** |  |  |
| RNA easy Kit | Qiagen | 74134 |
| rRNA Depletion Kit | New England Biolabs | E63105 |
| Ultra II RNA library prep kit | New England Biolabs | E7770G |
| Multiplex Oligos for Illumina | New England Biolabs | E7335G |
| ***Deposited Data*** |  |  |
| Hi-C, RNA-Seq and ATAC-Seq from A375 and MDA-MB-231 cell lines | This study | GEO:GSE143678 |
| ***Experimental Models: Cell Lines*** |  |  |
| A375 | ATCC | CRL-1619 |
| MDA-MB-231 | ATCC | CRM-HTB-26 |
| ***Transwell Migration Experiments*** |  |  |
| Fibronectin | Corning | 354008 |
| Transwell Filters (12 μm) | VWR | 10769-224 |
| Transwell Filters (5 μm) | VWR | 10769-236 |
| ***Software and Algorithms*** |  |  |
| Bowtie2 | (Langmead & Salzberg, 2012) | http://bowtie-bio.sourceforge.net/bowtie2/index.shtml |
| C-World | Job Dekker Lab | https://github.com/dekkerlab/cworld-dekker |
| STAR | (Dobin *et al*, 2013) | https://github.com/alexdobin/STAR |
| MACS2 | (Zhang *et al*, 2008) | https://github.com/taoliu/MACS |
| HTSeq | (Anders *et al*, 2015) | https://htseq.readthedocs.io/ |
| DESeq2 | (Love *et al*, 2014) | https://bioconductor.org/packages/release/bioc/html/DESEQ2.html |
| Bedtools | (Quinlan & Hall, 2010) | http://bedtools.readthedocs.io/en/latest/ |
| Samtools | (Li *et al*, 2009) | http://samtools.sourceforge.net/ |
| BBTools |  | https://github.com/kbaseapps/BBTools |
| CNV estimation | (Servant *et al.*, 2018) | https://github.com/nservant/cancer-hic-norm |
| Agent Based Modeling | Modified from https://github.com/zcolburn/cell-sheet-abm | https://github.com/rpmccordlab |

## CONTACT FOR REAGENT AND RESOURCE SHARING

Further information and requests for resources and reagents should be directed to and will be fulfilled by the Lead Contact, Rachel Patton McCord.

## EXPERIMENTAL MODEL AND SUBJECT DETAILS

### Cell Lines and Cell Culture

A375 and MDA-MB-231 cells were obtained from ATCC (CRL-1619 and HTB-26, respectively). Cells were verified to be negative for mycoplasma and were grown using complete DMEM medium (Corning – 10-013-CV; 10% FBS, 1% Pen-Strep, 1% L-Glutamine) at 37°C supplied with 5% CO_2_.

## METHOD DETAILS

### Sequential Transwell Migration

For the sequential migration, we used Transwell filters with 12 μm (VWR-10769-224) and 5 μm pore sizes (VWR-10769-236). Briefly, the bottom of the filters were coated with 40 μL of 10 μg/mL fibronectin for ∼ 45 minutes. 24 well plates were prepared for Transwell migration assay adding 500 μL of 1x DMEM (Corning) with full supplements per well. A375 cells were detached from culture dishes at 80-90% confluency and aliquoted to 100,000 cells per 100 μL of 1xDMEM. Each Transwell was placed into its corresponding well of the 24 well plate and 100 μL of the cell suspension was added to the top of each filter. Cells were incubated at 37°C, 5% CO_2_ and allowed to migrate for 24 hours. After the 24-hour incubation, migration efficiency was quantified as follows. First, freely floating cells were removed from the top of the filter (unmigrated; “Top” cells) and from the well beneath the filter (migrated “Bottom” cells) and placed in two separate tubes. Then, 400 μL of trypsin was added into the bottom chamber of the 24 well plate and 200 μL of trypsin was added into the top chamber to detach any remaining attached cells. Recovered cells after trypsinization were added to the unmigrated or migrated tubes, accordingly. Cells were spun down (1000 rpm, 5min) and counted (using trypan blue) to calculate % migration such as: #bottom/(#top+#bottom). Additionally, a small aliquot of cells was saved for immunofluorescence (IF). The rest of the cells were seeded into wells of a 24-well plate to expand. When cells reached 80-90% confluency, another Transwell migration was performed (R2). A375 Top and Bottom cells were detached and counted. Two Transwell filters were prepared as previously described (one for Top and one for Bottom). Then 100,000 cells suspended in 100 μL of 1xDMEM without supplements were seeded into the Transwell filters. After 24-hour incubation at 37°C and 5% CO_2_, cells were trypsinized as previously described to quantify migration efficiency. Only the cells that were always on top (Top of Top) and the cells that were always on the bottom (Bottom of Bottom) were saved and grown for further sequential rounds of migration. This process was repeated for 10 and 20 rounds of migration which lead to the generation of A375-Top5, Bottom-5, Top20-5, Bottom20-5, Top-12 and Bottom-12, A375-12-M10, A375-5-NM20 and A375-5-M20.

### 2D Single Cell migration

#### Live cell imaging

To track single cell movement in 2D, A375-Control, Top-5 and Bottom-5 cells were seeded on wells of a 6-well plate at a 30,000 cells/well density. After cells were attached, live cell imaging was performed using the EVOS FL Auto microscope at 40X magnification. Images were acquired every 10 minutes for a 24-hour time period.

#### Cell morphology analysis

Parameters that describe cell morphology such as solidity and aspect ratio were quantified using the Shape Descriptors plugin in ImageJ.

### Immunocytochemistry

#### IF staining

Approximately 50,000 – 100,000 cells for each sub-population of A375 were seeded into either poly-D-lysine treated coverslips or 35-mm coverslip bottom dishes. Cells were allowed to attach overnight and then crosslinked with 4% formaldehyde for 10 min followed by three, 5-minute washes with PBS. After washing, cells were permeabilized with permeabilization buffer (10% goat serum, 0.5% Triton in PBS) for 1 hour at room temperature. After the incubation, primary antibodies (Lamin A – sc-376248, Lamin B1 – ab133741, H3K9me3 – ab176916), diluted in antibody dilution buffer (5% Goat serum, 0.25% Triton in PBS) were added and incubated overnight at 4°C. After primary antibody incubation, cells were washed 3 times with PBS for 5 minutes for each wash. Cells were then incubated with secondary antibodies (Alexa Fluor 594 and Alexa Fluor 488) per manufacturer’s directions (2 drops of secondary antibody/1mL of PBS – R37117 and R37120) for 30 minutes at room temperature. After secondary antibody incubation, cells were washed 3 times with PBS and sealed using mounting media with DAPI. Slides were treated in the mounting media for 24 hours before imaging.

Cells were imaged using a Leica Sp8 Confocal microscope was equipped with a 63x oil immersion objective.

#### Nuclear morphology analysis

To analyze the diameter (minor axis), aspect ratio and standard deviation of maximum projected nuclei, Image J Shape Descriptor plugin was used. For morphological analysis of 3D reconstructed nuclei, Image J Nucleus J plugin was used with default parameters. Lamin A/C coefficient of variance was quantified as the ratio of Lamin A/C standard deviation to Lamin A/C mean intensity.

#### Radial distribution of Lamin A/C and H3K9me3

Lamin A/C and H3K9me3 radial distribution was quantified using Measure Object Intensity Distribution module in Cell Profiler. The maximum projected nuclei were separated into four equidistant bins. Mean fractional intensities for each bin (4 total bins; starting from innermost (bin1) to outermost (bin4) radial position) was plotted.

#### Line scan analysis

To quantify the distribution of Lamin A/C and H3K9me3 in the nucleus we also employed line scan analysis of nuclear central slices in Control, Top-5 and Bottom-5 cells. Line scans of same length were drawn across major and minor axis of nuclei using Leica SP8 Image Analysis software and intensity distribution was recorded for each nucleus. We then created bins of equal size ranging from 0-1 and normalized the distance for all nuclei. The line scan data from all the nuclei was aggregated by taking the mean of intensity values for each bin window.

### Metaphase Spread and Chromosome Staining (mFISH)

A375 Control and Bottom-5 cells were grown to 80% confluency in a T-25 flask. Mitotic cells were enriched for by adding 100ng/mL colcemid (KaryoMAX) directly to culture media for 6 hours. Cells were then lifted with Trypsin and centrifuged at 200xg. Supernatant was removed and pellet dislodged by flicking. Cells were swelled by adding pre-warmed (37 °C) 0.075 M KCl dropwise while slowly vortexing and then placed in 37 °C water bath for 6 minutes. 1 mL of Carnoy fixative (3:1 of methanol to acetic acid) was added to cell pellets and gently inverted to mix. Cells were then spun down at 200xg, and the supernatant discarded. Cells were then resuspended in 5mL of cold (4°C) Carnoy fixative and again spun at 200xg followed by supernatant removal. Cells then were resuspended in 250µL of cold (4 °C) Carnoy fixative. A 12µL sample was dropped from 12 inches onto a cover slip to induce nucleus rupture and spread metaphase chromosomes. Chromosomes were then stained by adding 2.25 µL of chr13 and chr5 FISH probes (Metasystems) to each slide. Slides were then placed in a 75 °C oven for 2 min to co-denature samples. Coverslips were mounted and sealed with rubber cement and slides were incubated at 37 °C overnight in a humified chamber. Coverslips were then gently removed, and slides washed in 0.4x SSC pre-warmed to 72 °C for 2 min. Slides were drained followed by a wash in 2xSSC/0.05 Tween-20 at RT for 30 sec. Slides were then rinsed with ddH_2_O and allowed to air-dry for 1 min. 2 drops of mounting media containing a DAPI stain were placed on each slide and sealed with a coverslip before being imaged on a Metasystems Metafer platform with a Zeiss epifluorescence microscope at 63x.

### Western Blot

A375 cells were trypsinized as previously described, and approximately 5 million cells were collected by centrifugation (1000 rcf, 5 min, room temperature). The cell pellets were resuspended in 300 ml of RIPA buffer (Thermo Scientific, I89900), supplemented with protease inhibitors/EDTA (GenDepot 50-101-5485), and phosphatase inhibitors (GenDepot 50-101-5488). Cells were then incubated on ice for 10 minutes. DNA was degraded by adding 6 μl of micrococcal nuclease (0.5 U/ml stock, Thermo Scientific, FEREN0181) and 12 μl of calcium chloride (100 mM stock) to each lysate, followed by incubation at 37°C for 15 min. Nuclease activity was inhibited by subsequent incubations at 68 °C for 15 min, and ice for 10 min. Protein samples were stored at −80 °C until use. Protein concentration was measured using the BCA Protein Assay Kit. Thermo (234225), according to the manufacturer’s instructions.

Denatured protein samples were resolved using a mini gel tank (Invitrogen, A25977) for 35 min at 200V, using 4-12% Bis-Tris Plus gels (Invitrogen, NW04120BOX) for all targets. Protein was transferred to low fluorescence PVDF membranes using the mini Bolt Blotting System (Invitrogen, B1000), along with system-specific reagents. Blotting was performed using the Odyssey TBS blocking system (Licor, 927-400000), according to the manufacturer’s protocol. Briefly, membranes were activated with methanol after transfer (1 min, RT), washed twice with milliQ water (5 min), twice with TBS (2 min), and blocked with Odyssey TBS blocking solution for 1 hour at RT. Primary antibodies were diluted in blocking buffer (containing 0.2% Tween), and incubated over night at 4°C as follows: H3K9me3 (1:1000, ab176916), LaminA/C (1:1000, sc-376248) and LaminB1 (1:1000, ab133741). Beta actin was used as a loading control, using the appropriate antibody (1:10,000; rabbit polyclonal PA1-16889, Thermo Fisher or 1:10,000; mouse monoclonal MA1-140, Thermo Fisher) alongside with the targets of interest. Detection was carried out using the following secondary antibodies: goat anti-rabbit (1:10,000; IRDye 680RD [92568071]; Licor) and goat anti-mouse (1:10,000, IRDye 800CW [95-32210]; Licor). Secondary antibodies were diluted in Odyssey TBS blocking buffer containing 0.2% Tween and 0.01% SDS. Membranes were incubated in secondary antibody for 1 hour at RT, followed by three washes (TBST, 5 min each).

Images of near infrared fluorescent signal were acquired using an Odyssey scanner (Licor) in both the red and green channels. Signal was quantified using the Image Studio software (Licor).

### Hi-C

#### Library Preparation

Hi-C libraries were prepared for each cell sub-population as previously described (Golloshi *et al*, 2018) including two biological duplicates for each condition. Briefly, 10 million cells were grown in T-75 cell culture flasks to 80-90% confluency and crosslinked with 1% formaldehyde for 10 min. Crosslinked cells were then suspended in lysis buffer to permeabilize the cell membrane and were Dounce homogenized. Chromatin was then digested in-nucleus overnight using DpnII restriction enzyme. Digested ends were filled in with biotin-dATP, and the blunt ends of interacting fragments were ligated together. DNA was then purified by phenol-chloroform extraction. For library preparation, the NEBNext Ultra II DNA Library prep kit (NEB) was used for libraries with size ranges from 200-400bp. End Prep, Adaptor Ligation, and PCR amplification reactions were carried out on bead bound DNA libraries.

For the clone migration experiments, Hi-C was performed using the Arima-HiC+ Kit from Arima Genomics following the protocol for Mammalian Cell Lines (A160134 v01) and preparing the libraries using Arima recommendations for the NEBNext Ultra II DNA library prep kit (protocol version A16041v01).

Sequencing was performed at the Oklahoma Medical Research Foundation Clinical Genomics facility or Azenta Genomics using the Illumina HiSeq 3000 platform with 75 bp paired end reads or a NovaSeq 6000 with 50 or 150 bp paired end reads. Sequencing reads were mapped to the human genome (hg19), filtered, and iteratively corrected as previously described (Imakaev *et al*., 2012) (https://github.com/dekkerlab/cMapping) or (for the clone Hi-C) with the same procedure but implemented by HiCPro (https://github.com/nservant/HiC-Pro). All Hi-C contact matrices were scaled to a sum of 1 million interactions to allow comparisons between conditions in downstream analysis. For library quality and mapping statistics see Table S1.

#### Compartment Analysis

Compartment analysis was performed by running principal component analysis using matrix2compartment.pl script in the cworld-dekker pipeline available on GitHub (https://github.com/dekkerlab/cworld-dekker). The PC1 value was then used to determine compartment identity for 100 and 250 kb binned matrices. We considered PC1 values greater than 0.01 to be A compartment and less than -0.01 to be B compartment for this analysis and determination of compartment switches.

#### Saddle Plots

Saddle plots were constructed to investigate changes in interaction frequency between or within compartments in highly migratory and non-migratory melanoma cells. Briefly, PCA analysis was performed on 100 kb binned matrices to assign compartment identity for each bin. For the analysis, interaction zScores, normalizing for interaction decay for genomic distance, were calculated according to the matrix2compartment.pl script in cworld-dekker. Zscore matrices were reordered based on compartment strength (from strongest B to strongest A using eigenvector values). Finally, matrices were smoothed at 500 kb bin size.

#### Interchromosomal Fraction (ICF)

Ratio of interchromosomal interactions relative to the total number of interactions for a given chromosome as previously described (Heinz *et al*, 2018).

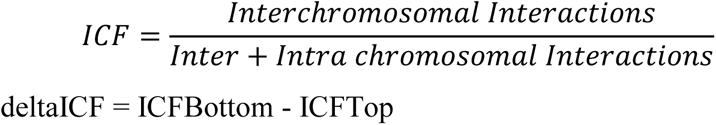

#### Copy Number Variation Estimation

CNV estimation was performed using the approach available from https://github.com/nservant/cancer-hic-norm (Servant *et al*., 2018)

#### Translocation Detection and Frequency Calculation

For the novel translocation visibly evident in 2.5 Mb binned contact maps for Bottom-5 constricted migrated cells, we further pinpointed the specific break site on each chromosome (chr5 and chr13) using raw valid pair counts within the region. First, we filtered all valid pair interactions between chr5 and chr13 to include only regions of chr13 between 38.3 and 57.5 Mb (a range of coordinates upstream of the translocation site, which all interact more than expected with chr5). Then, we plotted a histogram of interaction frequencies between each locus along chr5 and this set of chr13 coordinates. We noted a sharp interaction drop within the region chr5:12.24-12.25 Mb, which represents the translocation breakpoint. We then reversed this process to determine the breakpoint along chr13.

To calculate the percent of chromosomes containing this chr5-13 translocation in each subpopulation, we first found the 2.5 Mb bin with the maximum raw Hi-C counts near the translocation site in Bottom-5 cells. We then calculated an average value for raw Hi-C counts between neighboring bins (one bin off diagonal) in cis on chr7. We then calculated the percent of chromosomes bearing the translocation in any given cell type as the ratio of raw interaction counts at the translocation bin over average interaction counts between neighbors in cis. Given the median triploid karyotype of A375 cells, we then multiplied this % of chromosomes by 3 to estimate a % of cells carrying the translocation.

We followed a similar process as above for other detected translocations in A375 and MDA-MB-231 cells. Briefly, we identified all apparent translocations in the Hi-C map by searching for stronger than expected interchromosomal interactions with sharp boundaries on one edge signifying a chromosome break. We then identified the 2.5 Mb bin in this region that had a local maximum contact frequency and used this value as the contact strength of the translocation. We then calculated the expected contacts between adjacent regions of the chromosome as the average diagonal contacts on both chromosomes involved in the translocation. The translocation bin contacts divided by the average diagonal contacts were reported as the Translocation Intensity, a value proportional to the number of cells that carry this translocation in the cell population.

#### Distance Decay Scaling Plots

Using 40 and 20 kb binned iteratively corrected and scaled contact matrices, we extracted contact frequencies between bins at each genomic distance, excluding the diagonal bin (zero distance). A loess fit was then used to find a smooth curve describing interaction decay vs. distance (using the matrix2loess.pl script in cworld-dekker). The interaction frequencies were then normalized to set the maximum value (loess fit interaction value for the minimum distance) for each dataset to 1 and then plotted on a log scale vs. log genomic distance.

### RNA-Seq

#### Library Preparation

For Control, Top-5 and Bottom-5 A375 cells, RNA was extracted, and the library was prepared in our laboratory. Briefly, RNA was extracted using Qiagen RNAeasy plus mini kit (Cat. No. 74134). Cells were lysed and homogenized, spun down in gDNA eliminator columns to remove any genomic DNA. All samples were washed with ethanol and the total RNA was eluted. The quality and quantity of the RNA were quantified using Agilent Bioanalyzer Nano RNA kit. All the libraries used were characterized by a RIN value between 9 and 10. rRNA was depleted using NEBNext rRNA Depletion Kit (Cat. No. E6310S). Total RNA was hybridized to the probe followed by RNaseH and DNase I digestion. RNA was then purified using AmpureXP beads from Beckman Coulter (Ref. A63881) at a 2.2x concentration of beads. RNA libraries were prepared using NebNext Ultra II RNA library prep Kit (Cat. No. E7770G). After rRNA depletion, RNA was fragmented (∼200 nt) and primed following first and second strand cDNA synthesis. The double stranded cDNA was then purified using AmpureXP beads at a 1.8X concentration. After purification, cDNA libraries were end prepped, ligated adapter and PCR amplified for 7 cycles to enrichment for adaptor ligated DNA using NebNext Multiplex Oligos for illumine (Cat. No. E7335G). cDNA libraries were then purified using AmpureXP beads at a 0.9X concentration and their quality was checked using Agilent Bioanalyzer high sensitivity DNA kit. cDNA samples were then sent to Oklahoma Medical Research Facility (OMRF) for high throughput sequencing.

For the rest of A375 libraries, Top20-5, Bottom20-5, Top-12 and Bottom-12, RNA was extracted using the Qiagen RNAeasy plus mini kit as described above. Total RNA was then sent for library preparation and sequencing at GeneWiz. For library quality and mapping statistics see Table S2.

#### RNA-Seq Data Analysis

Quality of reads was checked using Fastqc and adapter sequences were trimmed using BBTools (https://github.com/kbaseapps/BBTools). Additionally, quality trimming of the reads was performed and any reads with a lower than 28 quality score were discarded. Reads were then aligned using STAR alignment (https://github.com/alexdobin/STAR) with an average alignment rate of 89%. The aligned reads were then sorted by genomic position and feature counts was performed using htseq 0.11.1(https://github.com/simon-anders/htseq). Differential expression of genes was determined by using DESeq through Galaxy.

### ATAC-Seq library preparation and analysis

ATAC-Seq libraries for Top-5 and Bottom-5 cells was prepared as previously described (Buenrostro *et al*, 2015). Libraries were generated using Ad1_noMX and Ad2.1-2.3 barcoded primers from (Buenrostro *et al*., 2015) and were amplified for a total of 12-13 cycles. The quality and quantity of the libraries were measured using Agilent Bioanalyzer High Sensitivity DNA kit. Sequencing was performed at Oklahoma Medical Research Facility using HiSeq3000 with 75 bp paired end reads. For library quality and mapping statistics see Table S3.

#### ATAC-Seq Data Analysis

Adapters and reads with a quality score less than 20 were trimmed from the final libraries using Skewer/0.2.2 (https://github.com/relipmoc/skewer). Trimmed reads were aligned to the hg19 genome using bowtie2 with default parameters (https://github.com/BenLangmead/bowtie2). Aligned reads were sorted, indexed and chrM reads were removed using samtools. Duplicated reads were removed using Picard (https://github.com/broadinstitute/picard/blob/master/src/main/java/picard/sam/markduplicates/MarkDuplicates.java). Finally, peaks were called using MACS2 (https://github.com/taoliu/MACS) with the following parameters: --nomodel --extsize 300 -g hs --keep-dup all.

### 3D Collagen Matrix

Experiments investigating the morphology and migration of A375 subpopulation of cells in 3D environment were performed using single cell suspension in 3D collagen matrices (A375-Control, A375-Top5 A375-Bottom10 generated from passage of A375 through 5 and 12 μm sized pores) as previously described (Denais *et al*., 2016). Briefly, 75,000 cells/condition were added into collagen matrices (Rat Tail, Corning 354236) with a density of 2.32 mg/ml. The pH of the collagen gels was normalized to a neutral pH and gels were incubated at 37^0^C for 30 min to gel. After the incubation, media with full supplements was supplied to the collagen gels. Cell were allowed to migrate through the collagen for 5 days before the cells were stained and imaged.

In timelapse images of cells embedded in collagen matrices, we found that about 5% of cells underwent apoptosis during the timecourse. We ensured that cells undergoing apoptosis were excluded from any analyses.

#### Immunofluorescence for 3D collagen matrices

Cells embedded on collagen were stained as previously described (Denais *et al*., 2016). Briefly, collagen matrices were fixed in 4% formaldehyde for 30 min at 37C. Collagen matrices were then washed with 1xPBS for 10 min for three times. Each collagen matrix was then incubated with blocking buffer (0.5% Triton, 10% Goat Serum in 1xPBS) overnight at 4C. After blocking, samples were then incubated with primary antibody (LaminA – sc376248) at a 1:500 dilution in 1xBlocking Buffer:1xPBS at 4C overnight. After incubation, aspirate primary antibody solution and wash three times with 1xPBS for 10 min each. Samples were then incubated in secondary antibody (2 drops/mL of PBS) for 1hr and room temperature. After secondary antibody incubation, samples were then washed with 1xPBS three times for 10 min each. Samples were incubated with DAPI (1μg/mL) for 30 min at RT. Collagen matrices were then washed three times with 1xPBS for 10 min each. Z-stacks of collagen matrices were taken with Leica SP8 confocal microscope at 40x water objective.

#### Image analysis for cells embedded in 3D collagen matrices

Aspect ratio and Lamin A/C standard deviation of maximum projected nuclei was quantified using ImageJ plugin Shape Descriptors. Tracking of cells migrating in 3D collagen matrices was performed using ImageJ Manual Tracking plugin. The resulting data was then used in Chemotaxis and Migration tool (Ibidi) to generate accumulated distance and velocity measurements.

### Agent Based Modeling of Constricted Migration

We model constricted cell migration using an on-lattice two-dimensional (2D) agent-based model (ABM) system. In this model, cells are represented by agents. The basic building block of this agent-based system is obtained from the collective cell migration ABM (collectiveABM), available on GitHub (https://github.com/zcolburn/cell-sheet-abm) and necessary changes are made to model cell transmigration. Most of the parameter values and the model working procedures of collectiveABM are kept untouched unless otherwise mentioned. Since the collectiveABM models a wound healing assay, we exclude the wound scratch section of the code from our model. We use two squared 2D grid spaces of size 20x20 to the agent-based system in place of a single 2D grid space: one to represent the top of the Transwell, where the cells are seeded at the beginning of each round of sequential migration, and another to represent the bottom of the Transwell, where the cells transmigrate from the top compartment. The beginning of each round of sequential migration model starts with 2000 cells seeded on the top compartment. The population of the cells is either considered to be homogenous or heterogeneous depending on the transmigration probability (Pmig). A total of 25 timepoint iterations are implemented to mimic the ∼24 hour duration of experimental migration rounds. At each timestep, each cell can choose to either transmigrate to the bottom, divide, die, or move horizontally. After each round of sequential migrations, the proportion of cells in the bottom and top compartment is measured, and the cells from the bottom or top are propagated to the next simulated round of migration. The directionality of the transmigration is only from top to bottom compartment and once the cells reach the bottom compartment, they stay there. The constricted migration of a cell is modeled by comparing its assigned Pmig with a uniform random number drawn from the range [0,1). If the random number is less than Pmig, then that cell will transmigrate to the bottom compartment from the top. In addition to having models with fixed Pmig values, we also designed agent-based model systems having variable Pmig : 1) the Pmig value of bottom cells are boosted by a function after each round and 2) each cell has different Pmig, drawn from a normal distribution and after each round, the mean and standard deviation of top or bottom compartment cells depending on the sequential Transwell migration experiment type will be used in next round’s normal distribution. The efficiency of migration is calculated at the end of each round by dividing the number of cells in the bottom compartment by total cells in the top and bottom compartments combined. For all the different test cases, three independent runs are performed by using different random number seeds.

## DATA AND SOFTWARE AVAILABILITY

All relevant data supporting the key findings of this study are available within the article and its Supplementary Information files or from the corresponding author on reasonable request. Hi-C, RNA-Seq, and ATAC-Seq sequence data and processed data for A375 cell line experiments are deposited under GEO: GSE143678. Software and code used for the analyses presented are available as public github packages, as described in the Methods section.

